# Cytokinin functions as an asymmetric and anti-gravitropic signal in lateral roots

**DOI:** 10.1101/572941

**Authors:** Sascha Waidmann, Michel Ruiz Rosquete, Maria Schöller, Heike Lindner, Therese LaRue, Elizabeth Sarkel, Ivan Petřík, Kai Dünser, Shanice Martopawiro, Rashmi Sasidharan, Ondrej Novak, Krzysztof Wabnik, José R. Dinneny, Jürgen Kleine-Vehn

## Abstract

Directional organ growth allows the plant root system to strategically cover its surroundings. Intercellular auxin transport is aligned with the gravity vector in the primary root tips, facilitating downward organ bending at the lower root flank. Here we show that cytokinin signaling functions as a lateral root specific anti-gravitropic component, promoting the radial distribution of the root system. We performed a genome-wide association study and revealed that signal peptide processing of Cytokinin Oxidase 2 (CKX2) affects its enzymatic activity and, thereby, determines the degradation of cytokinins in natural *Arabidopsis thaliana* accessions. Cytokinin signaling interferes with growth at the upper lateral root flank and thereby prevents downward bending. Our interdisciplinary approach revealed that two phytohormonal cues at opposite organ flanks counterbalance each other’s negative impact on growth, suppressing organ growth towards gravity and allow for radial expansion of the root system.

## Introduction

Root architectural traits define plant performance and yield (Uga et al., 2013). The radial spreading of the root system depends on the directional growth of primary and secondary roots. The phytohormone auxin plays a central role in aligning root organ growth towards gravity (Su et al., 2017). In the root tip, columella cells perceive changes in gravity via statolith sedimentation (Leitz et al., 2009). The relative change in statolith positioning triggers a partial polarization of redundant PIN3, PIN4 and PIN7 auxin efflux carriers towards this side, leading to enhanced auxin transport along the gravity vector (Friml et al., 2002; Kleine-Vehn et al., 2010). The asymmetric distribution of auxin eventually reduces cellular elongation rates at the lower root flank, which consequently leads to differential growth within the organ and bending towards gravity (Friml et al., 2003; Rosquete et al., 2013; Ruiz Rosquete et al., 2018).

Lateral roots (LRs) substantially differ from main roots, establishing a distinct gravitropic set point angle (GSA) (Digby and Firn, 1995). The divergent developmental programs of lateral and main (primary) roots allow the root system to strategically cover the surrounding substrate. In *Arabidopsis*, LRs emerge from the main root at a 90° angle (stage I LRs) and afterwards display maturation of gravity sensing cells, as well as the *de-novo* formation of an elongation zone (Rosquete et al., 2013). Transient expression of PIN3 in columella cells temporally defines asymmetric auxin distribution and differential elongation rates in stage II LRs (Guyomarc’h et al., 2012; Rosquete et al., 2013). This developmental stage lasts 8-9 hours and is characterized by asymmetric growth towards gravity at a slower rate than in primary roots (Rosquete et al., 2013; Schöller et al., 2018). During this phase of development, the primary GSA of LRs is established. The subsequent repression of PIN3 in columella cells of stage III LRs coincides with symmetric elongation, maintaining this primary GSA (Rosquete et al., 2013). Notably, the de-repression of PIN3 and PIN4 in columella cells of older stage III LRs does not correlate with additional bending to gravity (Ruiz Rosquete et al., 2018). This finding illustrates that the primary GSA is developmentally maintained, determining an important root architectural trait. Moreover, a stage III LR will return to its initial GSA if it is reoriented relative to the gravity vector (Mullen and Hangarter, 2003; Rosquete et al., 2013; Roychoudhry et al., 2013). Accordingly, the partial suppression of a full gravitropic response in recently emerged LRs is critical for establishing the primary growth direction of LRs, which importantly contributes to the root system architecture.

Despite the apparent importance of directional LR growth for radial exploration of the root system, the underlying suppressive mechanisms are largely unexplored. Using genetic, physiological, computational, biochemical, and cell biological approaches, we reveal that two opposing hormonal cues at the lower and upper lateral root flank counterbalance each other and set directional LR growth.

## Angular lateral root growth displays substantial natural variation in *Arabidopsis thaliana*

To examine the natural diversity in radial root growth, we screened 210 sequenced *Arabidopsis* accessions (Figure 1A, Table S1) and quantified their primary GSA of LRs. When grown *in vitro* on the surface of the growth medium, we observed extensive natural variation for the mean GSA values, detecting a deviation of about 40° between most extreme accessions (Figure 1B).

**Figure 1.**
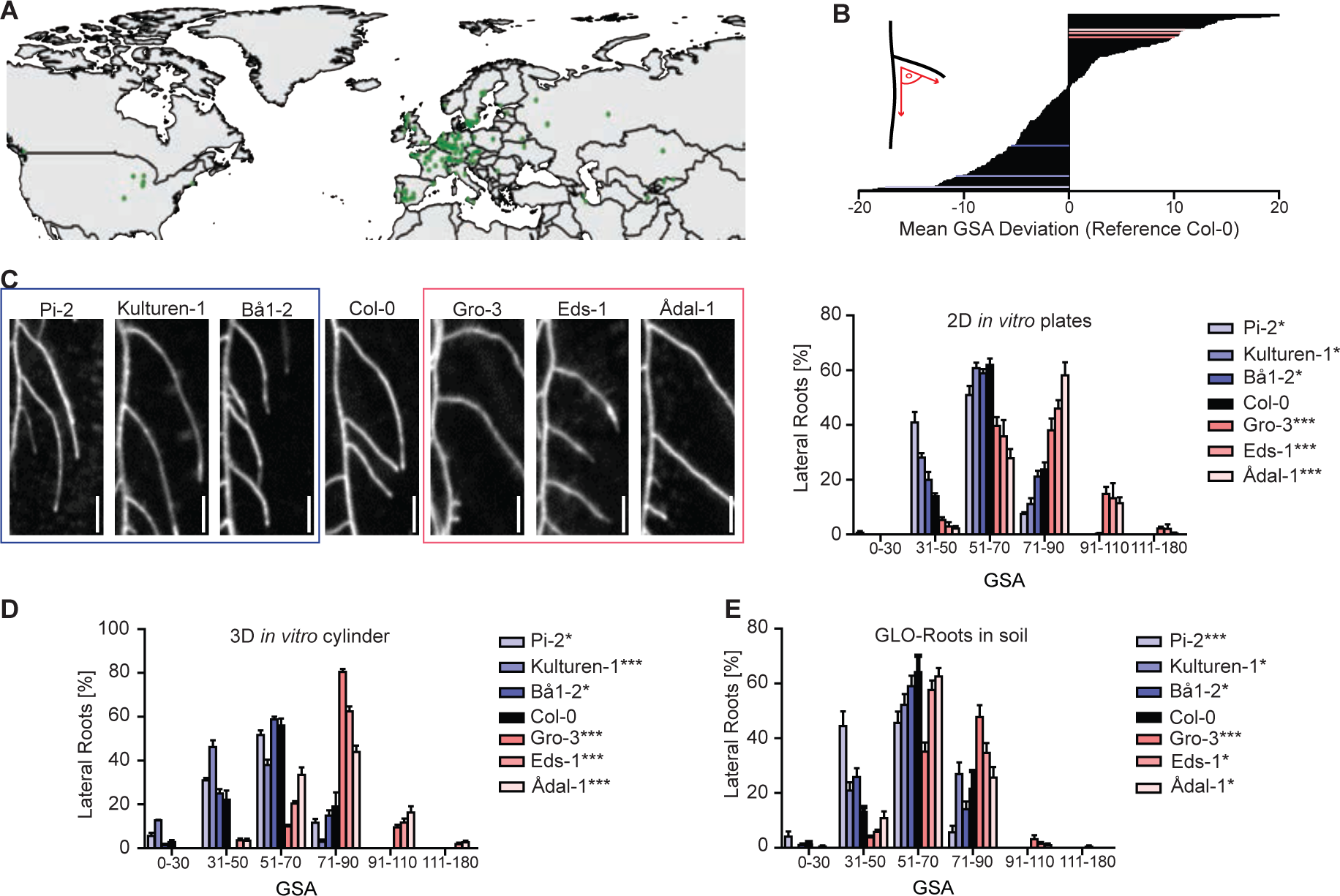
Natural variation of the primary GSA of lateral roots in *Arabidopsis thaliana*. (A) Geographical distribution of natural *Arabidopsis thaliana* accessions used in this study. (B) Mean gravitropic set point angle (GSA) values are normalized to reference accession Col-0. Three representative hyper- (blue colours) and hypo-responsive (red colours) accessions were selected for further analysis. (C) Representative images and GSA distributions of hyper- and hypo-responsive accessions grown on 2D agar plates. n = 5 plates (16 seedlings with 100-160 LRs per plate), Scale bars, 20 mm. (D) GSA distribution of hyper- and hypo-responsive accessions grown in 3D agar cylinders. n = 5 cylinders (20-40 LRs per cylinder). (E) GSA distribution of hyper- and hypo-responsive accessions grown in soil. n = 5-10 plants (20-40 LRs per plant). (C)-(D) Kolmogorov-Smirnov test P-values: * P < 0.05, ** P < 0.01, *** P < 0.001 (compared to Col-0). Mean ± SEM. Experiments were repeated at least three times.

Because the *in vitro* approach allowed only two-dimensional analysis of root growth, we further assessed angular growth of LRs in three dimensional and soil systems. For this purpose, we studied a subset (depicted by red and blue lines in Figure 1B) of hyper- and hypo-responsive accessions in greater detail (Figure 1C). To allow three-dimensional root expansion *in vitro*, we grew this subset of accessions in growth medium-filled cylinders (Ruiz Rosquete et al., 2018) (Figure S1A). In addition, we used the GLO-Roots system (Rellán-Álvarez et al., 2015), which is a luciferase (LUC)-based imaging platform to visualize root systems in a soil environment (Figure S1B). Accordingly, we transformed the same subset of accessions with *pUBQ:LUC2o,* ubiquitously driving LUC expression. In the Col-0 reference accession, about 60% of emerged LRs displayed a GSA between 51° and 70° in all three growth conditions (Figure 1C-E). In all growth systems, hypo- and hyper-responsive accessions displayed a pronounced shift towards higher (71°-90° and 91°-110°) and lower (31°- 50°) angle categories, respectively (Figure 1C-E). This suggests that our two-dimensional, *in vitro* screen is highly suitable to identify natural accessions with diverging GSA values of their root systems.

## Genome wide association study reveals a link between cytokinin metabolism and angular growth of lateral roots

Next, we sought to identify molecular players involved in the LR trait of our interest. To achieve this, we used our quantitative data on primary GSA of LRs and conducted a genome-wide association study (GWAS) (Seren et al., 2012). We identified several chromosomal regions, displaying associations with our trait (Figure 2A). A prominent peak at chromosome 2 drew our attention to a thymine (T)/guanine (G) single-nucleotide polymorphism (SNP) located in the *CYTOKININ OXIDASE2* (*CKX2*) gene (position 8,447,233) (Figure 2B). Importantly, the minor G allele, showing a frequency of 19.5% in all sequenced and 32.7% in our set of accessions, was associated with increased GSA values, reflecting more perpendicular LR growth to gravity (Figure 2C). Notably, linkage disequilibrium analysis showed that adjacent SNPs display pronounced non-random association with our SNP of interest (Figure S2), suggesting that the *CKX2* gene could be linked to variations in angular growth of LRs.

**Figure 2.**
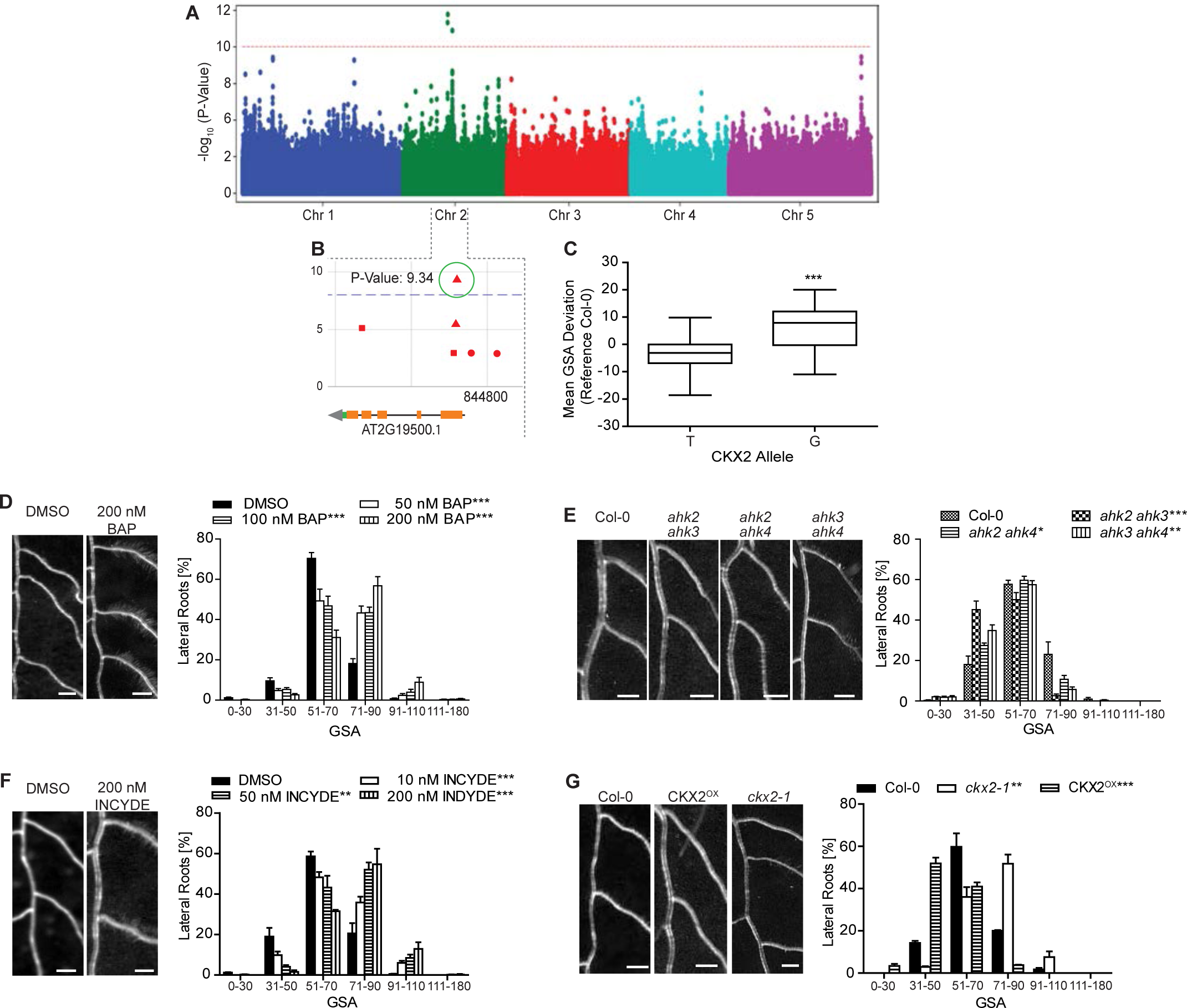
Genome-wide association study (GWAS) on gravitropic set point angle (GSA). (A) Manhattan plot of GWAS results. The dotted horizontal line indicates a significance level of 0.1 after Bonferroni correction for multiple testing. (B) Magnification of the peak region on chromosome 2. A highly significant SNP was located at position 8,447,233 in the coding region of *CKX2*. (C) Mean GSA of T and G allele of CKX2. Horizontal lines show the medians; box limits indicate the 25th and 75th percentiles; whiskers extend to the min and max values. Student’s t-test p-value: *** P < 0.001. (D)-(G) Representative images and GSA distributions of untreated and 6- Benzylaminopurin (BAP)-treated Col-0 wild type (D), Col-0 wild type*, ahk2 ahk3*, *ahk2 ahk4* and *ahk3 ahk4* (E), untreated and INCYDE-treated Col-0 wild type (F), Col-0 wild type, *ckx2-1* and CKX2^OX^ seedlings (G). Kolmogorov-Smirnov test P-values: * P < 0.05, ** P < 0.01, *** P < 0.001 (compared to DMSO solvent or Col-0 wild type control). Mean ± SEM, n = 5 plates (16 seedlings with 100-160 LRs per plate). Scale bars, 2 mm. (D)-(G) Experiments were repeated at least three times.

CKX enzymes are responsible for the irreversible degradation of cytokinins (CKs) via the oxidative cleavage of their side chain (Schmülling et al., 2003). Indeed, CK metabolism was affected in *ckx2-1 (ckx2* in *Col-0* background) mutant roots (Figure S3A-E), suggesting that CKX2-dependent metabolism of CKs may contribute to GSA establishment in lateral roots. To test whether this class of phytohormones may regulate angular growth of LRs, we initially transferred 7-day old seedlings of the reference accession *Col-0* to medium supplemented with CKs. We observed a concentration-dependent increase in GSA values of LRs emerging in presence of active CKs, such as 6-Benzylaminopurin (BAP) (Figure 2D), trans-zeatin (tZ) and isopentenyladenine (iP) (Figure S3F and G). Conversely, CK receptor mutants showed accelerated bending of LRs and accordingly decreased GSA values (Figure 2E). These data suggest that cytokinin signaling interferes with downward bending of emerged LRs. Notably, emerging LRs of winter oilseed rape also displayed reduced bending of LRs when treated with BAP (Figure S3H), suggesting that the effect of CK on directional LR growth is likely to be conserved.

To further assess the importance of CKX2 in GSA establishment, we disrupted CKX activity in the reference accession *Col-0*. Treatments with the CKX inhibitor INCYDE (Zatloukal et al., 2008) phenocopied the *ckx2* loss-of-function mutant, both displaying more horizontal LRs when compared to its respective controls (Figure 2F, G). On the other hand, *CKX2* overexpressing (OX) plants showed accelerated bending of LRs, phenocopying the CK receptor mutants (Figure 2G). These data suggest that CK signaling defines directional lateral root growth by reducing LR bending after emergence.

## Cytokinin Response Factors define angular growth of lateral roots

Our data indicates that CK signaling impacts the angular growth of emerged LRs. Therefore, we assessed whether CK-dependent transcription factors indeed have an impact on LR growth in the reference accession *Col-0*. It has been previously shown that CK signaling initiates transcriptional changes via the *Arabidopsis* response factors (ARRs) (Skylar et al., 2010) and the cytokinin response factors (CRFs) (Raines et al., 2015; Šimášková et al., 2015). According to available organ specific microarray data (Brady et al., 2007), *ARR3* and *ARR4* as well as *CRF2* and *CRF3* (Figure S4A-B) were strongly upregulated in older LRs. However, we did not detect any expression of *ARR3* and *ARR4* in young stage II LRs, using promoter GUS reporter lines for *ARR3* and *ARR4* (*pARR3:GUS/pARR4:GUS;* Figure S4C)). Moreover, angular growth of LRs was not altered in *arr3* or *arr4* mutants (Figure S4D). On the other hand, we confirmed expression of *CRF2* and *CRF3* in the early stages of LR development (Figure 3A and S4E). *pCRF2:GFP-GUS* was ubiquitously expressed in young LRs, while *pCRF3:GFP-GUS* was preferentially expressed in cortical and epidermal cell files (Figure 3A and S4E). Notably, compared to emerged laterals, the main root displayed much weaker *CRF2* and *CRF3* expression (Figure S4F), suggesting that *CRF2* and *CRF3* are particularly highly expressed in young LRs.

**Figure 3.**
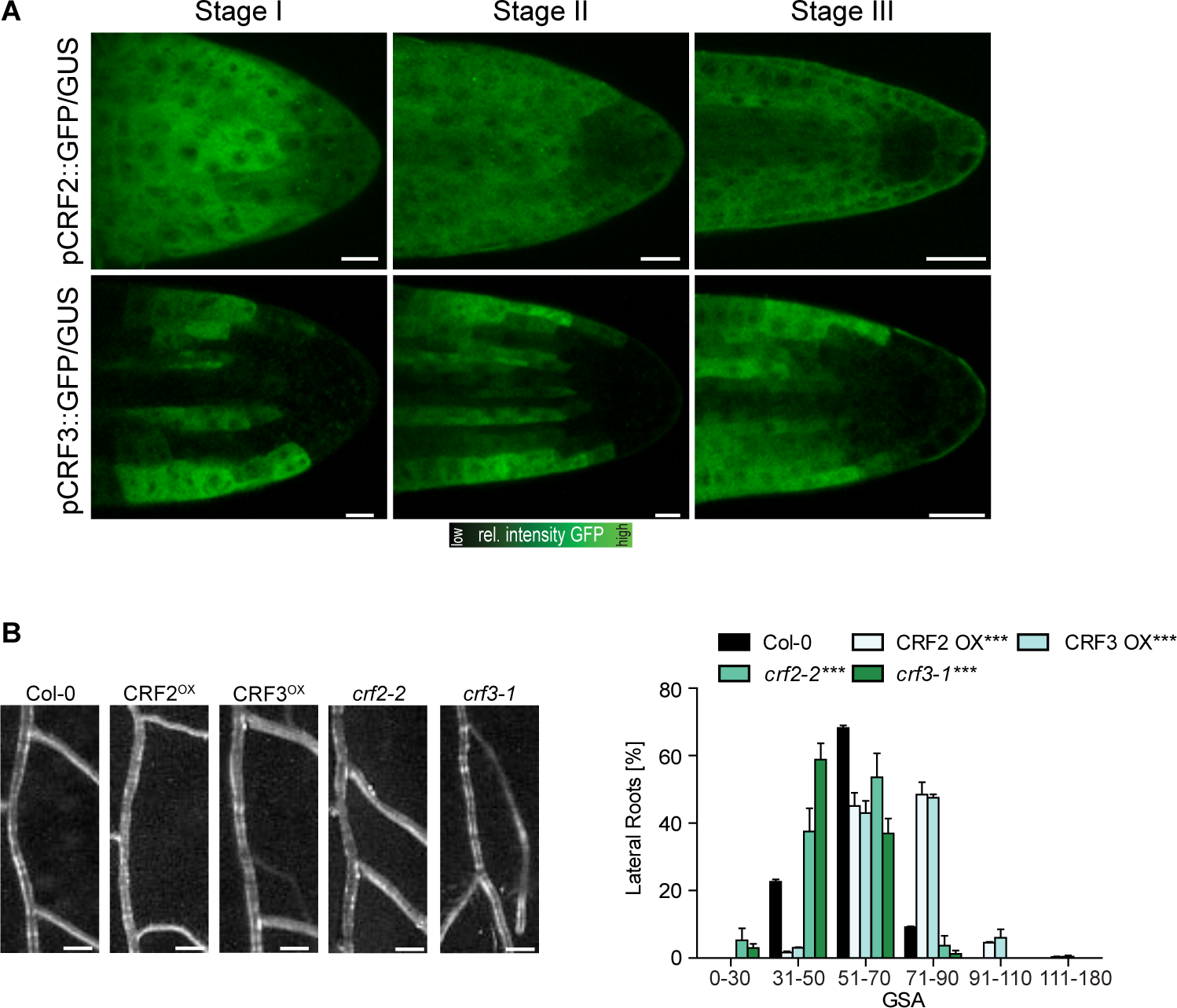
Characterization of Cytokinin Response Factors (CRFs) in lateral roots. (A) Representative images of pCRF2::GFP/GUS and pCRF3::GFP/GUS in stage I – III LRs. Scale bar, 25 µM. (B) Representative images and GSA distribution of Col-0 wild type, *crf* mutants and CRF^OX^ lines. Kolmogorov-Smirnov test P-values: *** P < 0.001 (compared to DMSO or Col-0, respectively). Mean ± SEM, n = 5 plates (16 seedlings with 80-100 LRs per plate). Scale bars, 2 mm. (A)-(B) Experiments were repeated at least three times.

In agreement with *CRF2* and *CRF3* expression in emerged LRs, loss-of-function alleles of *crf2* and *crf3* displayed enhanced bending of LRs (Figure 3B and S4G). Conversely, we found that ubiquitous overexpression of either CRF2 or CRF3 led to more horizontal LRs (Figure 3B).

This set of data confirms that cytokinin signaling, utilizing transcription factors such as CRF2 and CRF3, regulates angular growth of LRs.

## Cytokinin signaling integrates environmental cues into angular growth of lateral roots

Our data supports a role for cytokinin signaling in modulating angular growth of LRs. To investigate whether cytokinin modulates angular LR growth in response to environmental cues, we examined whether the primary GSA of Arabidopsis accessions is linked to geographic origins. Intriguingly, accessions with the largest GSA values predominantly originated in Nordic (above 58°N) regions (Figure 4A). In addition, the above described minor G allele of *CKX2,* phenocopying the *ckx2* loss of function (in *Col-0*), was notably the most prevalent allele in the north of Sweden (Figure 4B). Previous work showed that *Arabidopsis* accessions in the north of Sweden are fully vernalized before snow fall (Duncan et al., 2015). In fact, the respective habitat in the north of Sweden is most of the year covered with snow (Figure S5A). Snowpack insulation capacity can protect plants from extreme temperatures, but may also restrict soil-atmosphere gas exchange, eventually leading to hypoxia in the soil (Martz et al., 2016). Additionally, rapid snowmelt in spring can lead to temporary soil flooding, which depletes soil oxygen and may restrict the amount of oxygen reaching the root tissues. Hypoxic conditions have been shown to induce bending in the primary root as a possible adaptive avoidance response (Eysholdt-Derzsó and Sauter, 2017). Therefore, we asked whether hypoxia conditions also modulate the bending of LRs. In contrast to the primary root response, we observed that hypoxic stress reduced bending in emerged LRs (Figure 4C), demonstrating distinct pathways to regulate root bending in primary and secondary roots. Interestingly, hypoxia stress for 4 hours was sufficient to increase GSA of subsequently emerged LRs in *Col-0* (Figure 4C), mimicking the *ckx2* loss of function phenotype. Furthermore, hypoxic stress did not further increase GSA in the *ckx2* mutant (Figure 4C), proposing that CK metabolism could mediate hypoxia-dependent repression of LR bending. Furthermore, the LRs of *ahk2 ahk4* cytokinin receptor mutants were insensitive to the hypoxia-induced repression of LR bending (Figure 4D). Accordingly, we conclude that cytokinin signaling integrates environmental signals, such as hypoxia, into GSA establishment of emerged LRs.

**Figure 4.**
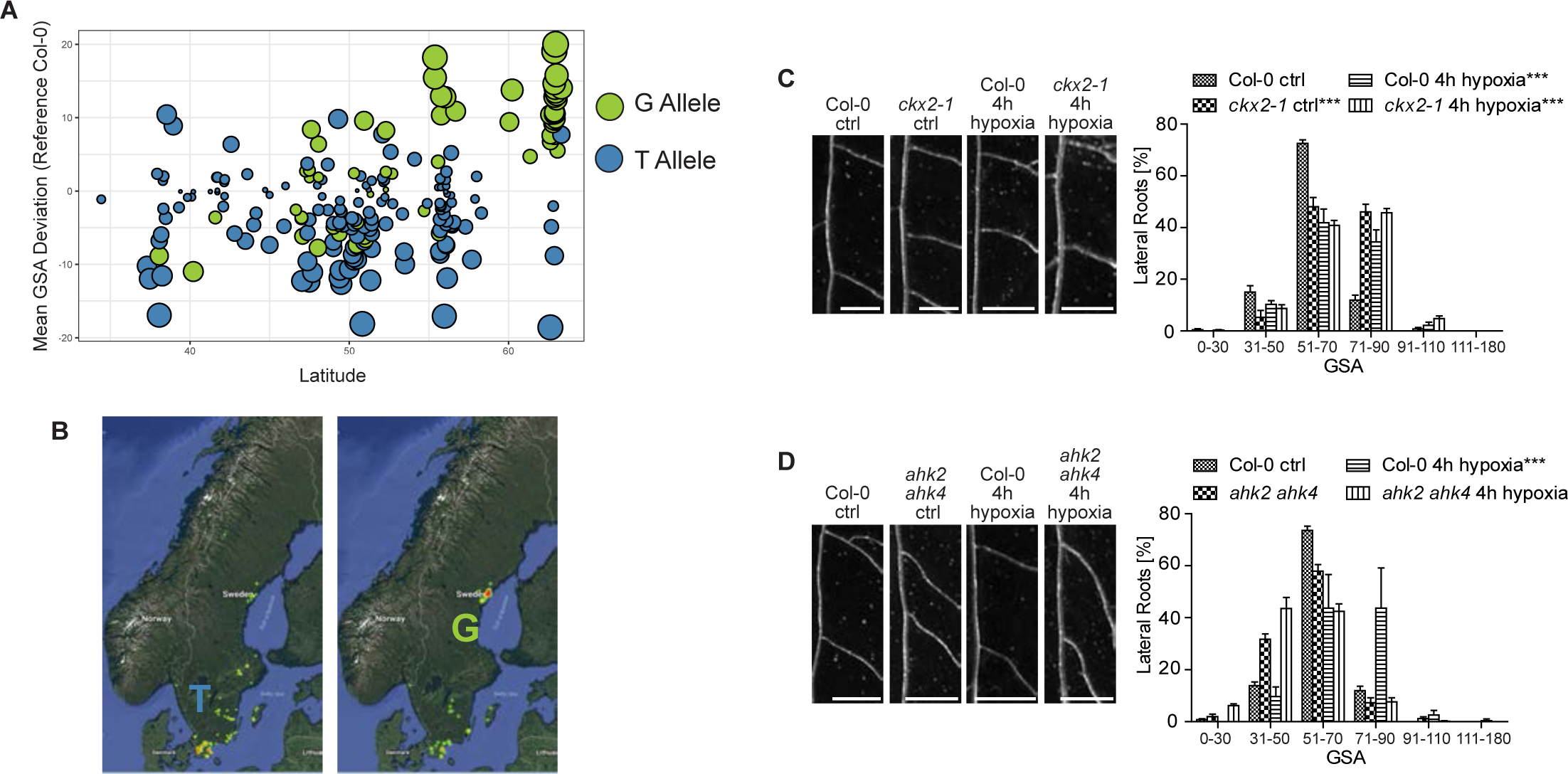
Cytokinin signalling integrates environmental signals into angular lateral root growth. (A) Comparison of the mean GSA distribution and its geographical (latitude) distribution of the phenotyped accessions. T and G allele of CKX2 are depicted in blue and green, respectively. (B) Relative geographical distribution (color coded by yellow (low) to red (high number)) of the T and G allele of CKX2 in all sequenced Swedish Arabidopsis accessions. (C)-(D) Representative images and GSA distributions of (C) Col-0 wild-type and *ckx2-1* or (D) Col-0 wild type and ahk2 ahk4 with and without hypoxia treatment for 4h. Scale bars, 2 mm. Kolmogorov-Smirnov test P-values: *** P < 0.001 (compared to DMSO solvent or Col-0 wild type control). Mean ± SEM., n = 4 plates (10 seedlings with 80- 100 LRs per plate). Experiments were repeated at least three times.

## Single base-pair variation in *CKX2* impacts on its *in-planta* activity

Our data proposes that variation in *CKX2* is linked to the control of radial root system expansion in natural *Arabidopsis* accessions. The previously mentioned G allele of *CKX2* is associated with higher GSA values (Figure 2C), which phenocopies the loss of *CKX2* function or increase in CK levels (Figure 2D). Accordingly, we next assessed whether the identified SNP affects the activity of CKX2. The underlying T to G mutation alters the first amino acid in the mature enzyme from an isoleucine (I) to a methionine (M). This mutation is situated just after the predicted cleavage site of a signal peptide (SP). The SP allows CKX2 to be inserted into the endoplasmic reticulum and to be subsequently secreted into the extracellular space (apoplast) (Schmülling et al., 2003). To assess potential trafficking or processing defects caused by the amino acid change (Samalova et al., 2006), we generated a ratiometric CKX2 reporter by fusing the Green Fluorescent Protein (GFP) and mScarlet to the N-terminal and C-terminal ends of CKX2, respectively. Fluorescent mScarlet signal of the non-mutated CKX2^I^ readily accumulated in the apoplast, suggesting that the fluorescent tags do not abolish processing and/or secretion of CKX2-mScarlet (Figure 5A and S5B). Ratiometric imaging of GFP and mScarlet revealed a higher degree of co-localization for mutated version CKX2^M^, suggesting reduced processing and/or secretion of CKX2^M^ when compared to CKX2^I^ (Figure 5B). To visualize the effect of the T to G mutation on SP cleavage, we N-terminally tagged CKX2 with GFP and subsequently expressed GFP^SP^CKX2^I^ and its respective mutated version GFP^SP^CKX2^M^ in tobacco. Western blot analysis revealed a decreased cleavage of GFP^SP^CKX2^M^ when compared to GFP^SP^CKX2^I^ (Figure 5C). Even though we cannot eliminate the possibility that N-terminal GFP may interfere with normal SP processing rates, the relative differences between the two assessed alleles suggests that the identified SNP impacts the SP cleavage in CKX2.

**Figure 5.**
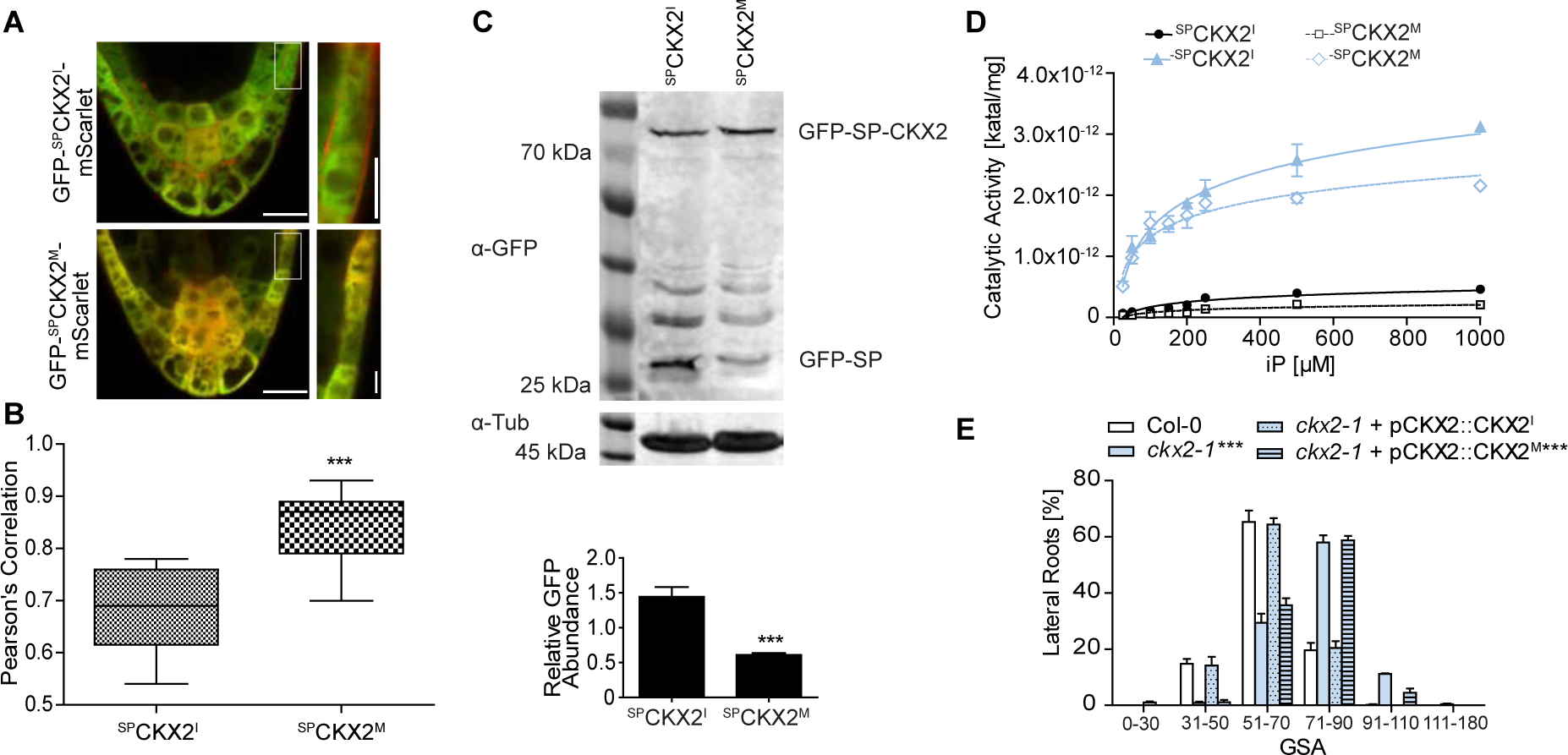
Signal Peptide processing is required for CKX2 activity. (A) Localization of GFP-^SP^CKX2^I^-mScarlet and GFP-^SP^CKX2^M^-mScarlet in stage II LRs. Scale bar, 25 µm and 10 µm, respectively. (B) Quantification of the co-localization of GFP and mScarlet signal using Pearson‘s correlation. Horizontal lines show the medians; box limits indicate the 25th and 75th percentiles; whiskers extend to the min and max values. Student’s t-test P-Value: *** P < 0.001, n = 10-15 individual LRs. (C) Immunoblot analysis and quantification of ^SP^CKX2^I^ and ^SP^CKX2^M^ expressed in *N. tabacum* leaves using anti-GFP antibody. Anti-tubulin antibody was used as loading control. The signal of GFP-SP was quantified and normalized to tubulin. Student’s t- test P-Value: *** P < 0.001. Mean ± SEM, n = 8 biological replicates. (D) Saturation curves of isopentenyladenine (iP) degradation by CKX2. Reactions were performed at pH 7.4 in McIlvaine buffer with 0.5 mM DCIP as electron acceptor (-●- ^SP^CKX2^I^, -□- ^SP^CKX2^M^, -▴- ^-SP^CKX2^I^, -◊- ^-SP^CKX2^M^). Mean ± SEM, n= 8. (E) GSA distributions of ckx2-1 was complemented by CKX2::CKX2I, but not by CKX2::CKX2M. Representative lines are shown. Kolmogorov-Smirnov test P-value: *** P < 0.001 (compared to Col-0). Mean ± SEM, n = 5 plates (16 seedlings with 100-160 LRs per plate). (A)-(E) Experiments were repeated at least three times.

The SP processing is an important determinant of the mature protein and, hence, we examined the enzymatic CKX2 activity in the presence and absence of the signal peptide. We expressed full length ^SP^CKX2^I^ and ^SP^CKX2^M^ as well as the SP-lacking counterparts ^-SP^CKX2^I^ and ^-SP^CKX2^M^ in *Escherichia coli* and measured their ability to oxidize CKs. Both SP-lacking forms ^-SP^CKX2^I^ and ^-SP^CKX2^M^ showed a 10-fold higher activity compared to the SP containing versions (Figure 5D). This *in vitro* data suggests that SP processing is required to ensure full enzymatic activity of CKX2.

Next, to assess whether the T to G mutation also affects CKX2 activity *in planta*, we expressed full length *pCKX2::CKX2^I^* and *pCKX2::CKX2^M^* encoding versions in the *ckx2* mutant background. As expected, the wild-type (Col-0) *CKX2^I^* was able to fully complement the *ckx2* mutant phenotype (Figure 5E and Figure S5C). In contrast, the mutated *CKX2^M^* version was not able to reverse the reduced LR bending of *ckx2* mutants (Figure 5E and S5C). Overall, our data suggests that the T to G mutation found in natural accessions renders *CKX2* to be largely non-functional *in planta* by disrupting its secretion and/or SP processing.

Thus, we conclude that variation in SP processing of CKX2 contributes to the natural variation of CK-dependent angular LR growth in *Arabidopsis*.

## CKX2 does not detectably interfere with auxin signaling in emerged lateral roots

Next, we investigated the cellular mechanism by which CKX2 activity modulates the primary GSA of LRs. We first inspected the spatial expression of *CKX2* to identify cells in which *CKX2* may directly regulate angular growth in LRs. *pCKX2::CKX2- mTurquoise* was weakly expressed in the tip of stage I LRs but showed increased expression in stage II and III LRs (Figure 6A). We confirmed that endogenous *CKX2* transcripts are strongly up-regulated in stage II and III LRs by examining expression in excised LR tissue using qPCR (Figure 6B). Notably, *pCKX2::CKX2-mTurquoise* was not detectable in the primary root tip (Figure 6B and S6A), proposing that CKX2 might specifically act in secondary root organs.

**Figure 6.**
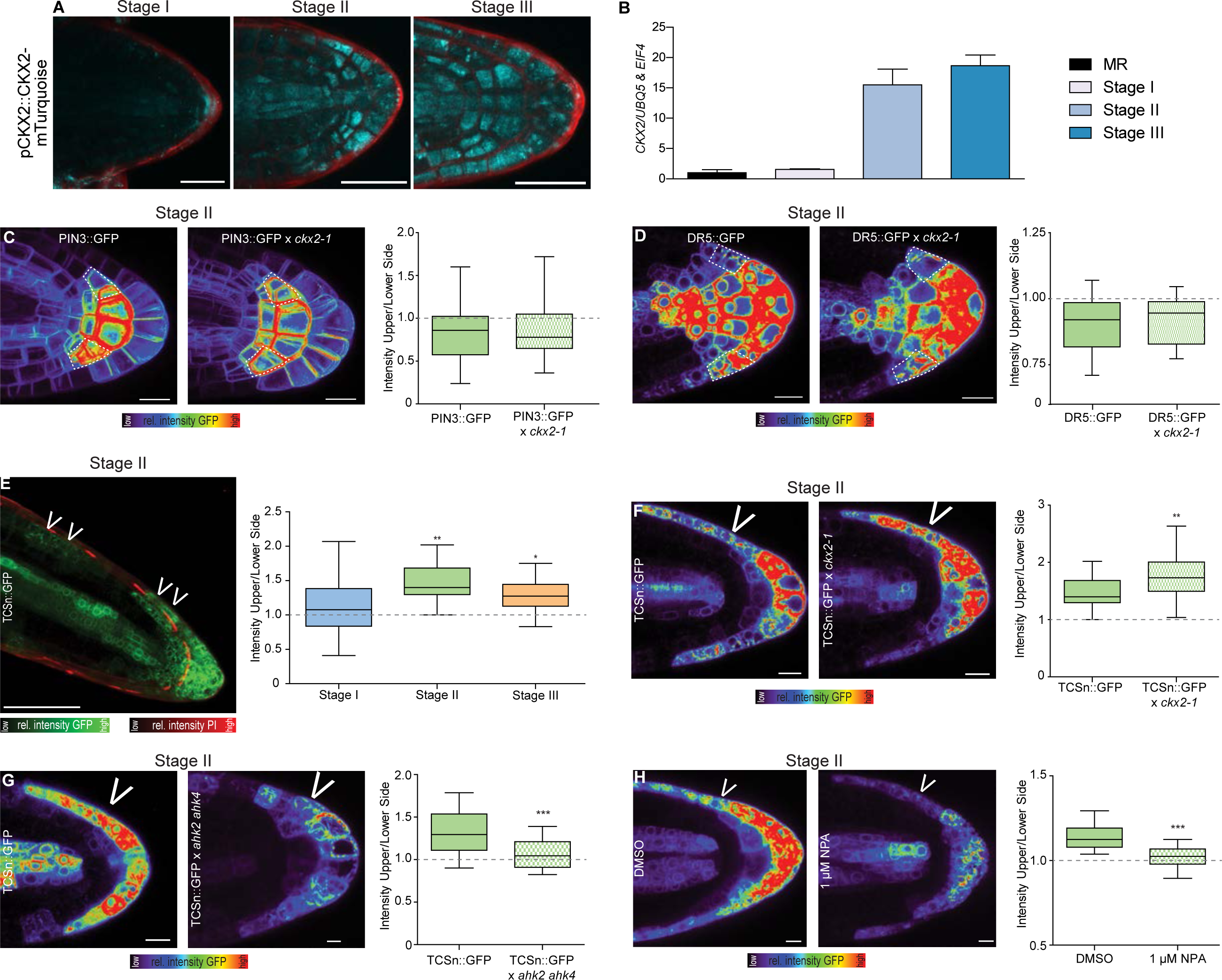
CKX2 modulates assymmetric cytokinin signalling in emerged lateral roots. (A) Representative images of pCKX2::CKX2-mTurquoise in stage I – III LRs. Propidium Iodide (PI) was used for counterstaining. Scale bar, 25 µm. (B) qPCR analysis detecting the levels of *CKX2* transcript in the root tip and LRs stage I-III normalized against UBQ5 and EIF4. Bars represent means ± SD, n = 3. (C)-(D) Representative images and signal quantification of stage II LRs of (C) PIN3::PIN3-GFP, and (D) DR5::GFP in Col-0 wild type and *ckx2-1* mutant background. Horizontal lines show the medians; box limits indicate the 25th and 75th percentiles; whiskers extend to the min and max values, n = 10-15 individual LRs. White dotted lines outline lateral root cap cells (facing the columella cells) for quantification. Scale bars, 10 µm. (E) Representative image (stage II) and quantification of TCSn::GFP in stages I – III LRs. PI was used for counterstaining. Scale bar, 50 µm. (F)-(G) Representative images and quantification of stage II LRs of (F) TCSn::GFP in wild type and *ckx2-1*, (G) TCSn::GFP in wild type and *ahk2 ahk4* or (D) after treatment with DMSO or 1 µM NPA for 24h. Scale bars, 10 µm. One-way ANOVA P-values: * P < 0.05, ** P < 0.01, *** P < 0.001. Horizontal lines show the medians; box limits indicate the 25th and 75th percentiles; whiskers extend to the min and max values, n = 10-15 individual LRs. (A)-(G) Experiments were repeated at least three times.

We next aimed to investigate how deviations in CKX2-dependent modulation of CK in LRs may modulate their directional growth. CKs signaling impairs PIN-dependent auxin transport in main roots as well as in lateral root primordia (Marhavý et al., 2011). We therefore assessed whether CKX2 activity regulates auxin transport in emerged LRs. Because PIN3 is the main regulator of asymmetric auxin redistribution in columella cells of emerged lateral roots (Rosquete et al., 2013), we initially assessed whether the *ckx2* mutant shows defective abundance or localization of functional pPIN3::PIN3-GFP in columella cells. At the time of GSA establishment (stage II LRs), PIN3-GFP abundance and asymmetry are not detectably altered from wild-type in *ckx2* mutants (Figure 6C and S6B). Next, we used the auxin responsive promoter DR5 fused to GFP and assess whether auxin signaling is affected in *ckx2* mutant LRs. In accordance with proper PIN3 localization, DR5 signal intensity in columella cells and asymmetric signal in the flanks was similar in *ckx2* mutant and wild type LRs (Figure 6D and S6C).

Overall, these data illustrate that auxin responses in gravitropic lateral roots are not detectably altered by CKX2, suggesting that CKX2 modulates angular growth by an alternative, CK-dependent mechanism in emerged LRs.

## Emerged lateral roots display asymmetric cytokinin signaling

Our data indicates that CK regulates angular LR growth. To further assess the mechanism by which CK modulates GSA establishment in developing LRs, we visualized the spatial distribution of CK signaling using the two-component signaling sensor (TCSn) transcriptionally fused to GFP (TCSn::GFP) (Liu and Müller, 2017). We observed increased CK signaling on the upper side of stage II LRs, coinciding with gravitropic bending (Figure 6E). This asymmetry declined in stage III LRs, which maintain the previously established GSA (Figure 6E and S6D). In agreement with the anticipated reduction in CK degradation, the magnitude of asymmetric CK signaling was increased in *cxk2* mutant LRs (Figure 6F and S6D). Conversely, asymmetric CK signaling was reduced in the CK receptor double mutant *ahk2 ahk4* (Figure 6G and S6E). These data propose that the increased magnitude of asymmetry in CK signaling across the root tip correlates with reduced LR bending towards gravity.

To determine whether asymmetric CK signaling regulates bending specifically in LRs, we examined the distribution of CK signaling in primary roots responding to gravity. Importantly, we did not observe asymmetric CK signaling in unstimulated or gravity-stimulated primary roots (Figure S6F-G). Accordingly, we conclude that asymmetric CK signaling is specific to LRs and thus contributes to the distinct establishment of primary GSA in LRs. Previous work proposed a hypothetical gravitropic offset component at the upper flank of LRs. This envisioned component was presumably sensitive to the inhibition of auxin transport (Roychoudhry et al., 2013). To assess if auxin transport similarly modulates the asymmetry of CK signaling in emerged LRs, we treated seedlings with the auxin transport inhibitor 1-N-Naphthylphthalamic Acid (NPA). Pharmacological interference with auxin transport indeed markedly decreased asymmetric CK signaling in stage II LRs, when compared to the DMSO solvent control (Figure 6H and S6H), suggesting that auxin transport indeed impacts asymmetric CK signaling in emerged LRs.

In summary, our data suggests that asymmetric CK signaling at the upper flank of LRs functions as an anti-gravitropic component in emerged LRs to promote radial root growth.

## CKX2 activity determines cellular elongation in emerged lateral roots

Light sheet-based live cell imaging has revealed that cells on the upper and lower flanks of emerged LRs show differential elongation for about 8-9 hours (Rosquete et al., 2013). During this developmental stage II, the cellular elongation rates at the upper epidermal layers is three-fold-increased compared to the lower flank (15µm/h versus 5µm/h) (Rosquete et al., 2013). To test if this difference can account for the primary GSA establishment, we used these quantitative growth parameters to construct a dynamic computational model of LR bending (Figure 7A-D and S7A-D). This model incorporates cellular mechanics to simulate cell elongation using stretchable strings as a manifestation of the cell wall elasticity and internal turgor pressure in the cell (see method section). The anisotropic growth is simulated by extending the resting length of the spring to account for 3-fold differences in the growth rates between upper and lower flanks. The resulting model predicts that the incorporation of measured elongation rates on the upper LR flank is able to realistically recapitulate LR bending angle of wild type plants, reaching an angle of about 62-63° within 8-9 hours (Figure 7B and Figure S7).

**Figure 7.**
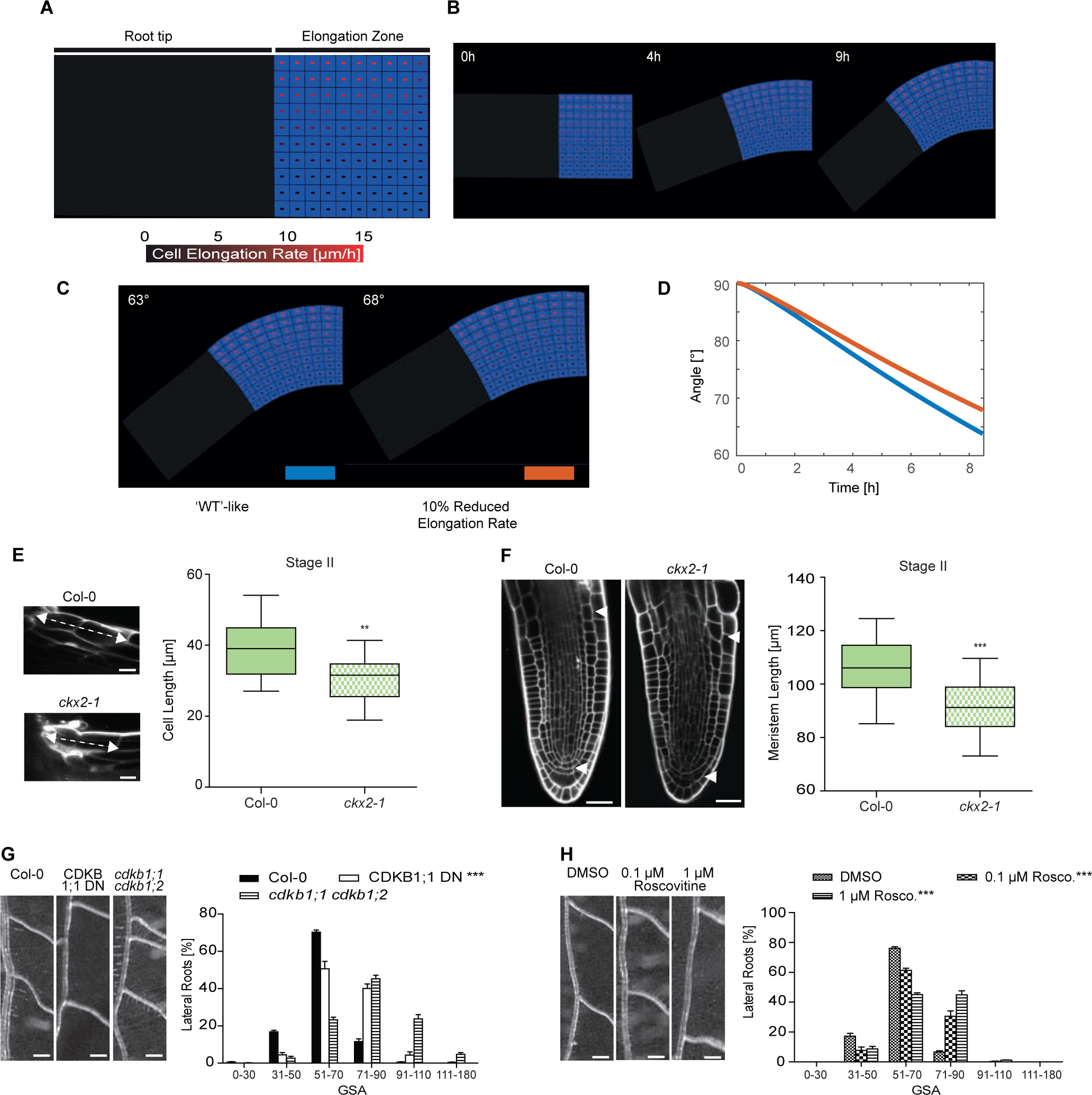
Cytokinin-dependent interference with cell cycle defines angular growth of lateral roots. (A) Sketch shows a simplified geometry of lateral root (LR), representing the root tip and the elongation zone. The cell elongation rate (visualized as red spot inside the cell) gradually increases from lower towards the upper side of the LR (∼ 3-fold) based on estimates derived from previous work (Rosquete et al., 2013). Bottom panel, color coding bar for cell elongation rates. (B) Time-lapse model simulations, considering 9 elongating cells establishing LR bending (63°) after approximately 9h. (C) left control panel (corresponds to (B)) is compared to right panel showing a 10% decrease in elongation rate on the upper root flank. Each simulation represents LR status after 8h of dynamic elongation. (D) Time evolution of set-point angle corresponding to (C), color of curves matches simulation with the color bar shown in (C). (E)-(F) Representative image and quantification of (E) first two elongated cells of lateral roots in stages II and (F) lateral root meristem. One-way ANOVA P-values: ** P < 0.01, *** P < 0.001. Horizontal lines show the medians; box limits indicate the 25th and 75th percentiles; whiskers extend to the min and max values, n = 10-15 individual LRs. Scale bar, 10 µm. (G)-(H) Representative images and GSA distributions of (D) Col-0 wild-type, CDKB1;1 DN (dominat negative) and cdkb1;1 cdkb1;2 or (E) Roscovitine treated Col-0 wild-type seedlings. Kolmogorov-Smirnov test P-values: ** P < 0.01, *** P < 0.001 (compared to Col-0). Mean ± SEM, n = 5 plates (16 seedlings with 60-80 LRs per plate). Scale bars, 2 mm. (E)-(H) Experiments were repeated at least three times.

Next, we experimentally assessed whether the loss of *CKX2* or CK application interferes with cell elongation in stage II LRs. In agreement with reduced LR bending, the *ckx2* loss-of-function mutant, as well as wild type plants treated with BAP, showed shorter cells at the upper flank of stage II LRs when compared to the respective controls (Figure 7E and S7E). Our previous work revealed that differential elongation is a major factor controlling LR bending (Rosquete et al., 2013). However, the loss of *CKX2* reduced cell elongation in average only by ten percent. To evaluate whether the measured reduction in cell length can realize the observed quantitative changes in LR bending, we reduced cellular elongation similarly by ten percent in our computational LR model. The model predicted that CKX2-dependent impact on cellular elongation increases the predicted GSA of LRs within nine hours from 63° to only 68° (Figure 7C-D). Thus, we conclude that the impact of CKX2 on cellular elongation cannot fully explain the observed reduction of LR bending in *ckx2* mutants.

## Cytokinin-dependent interference with cell division rates defines angular growth of lateral roots

In primary roots, CK reduces not only cellular elongation, but also cell proliferation by distinct mechanisms (Ruzicka et al., 2007; Street et al., 2015). Moreover, our computational model predicts that the rate of LR bending could be restricted by the number of cells (Figure S7A-B). Thus, we tested if CK might also affect the meristem of LRs. The stage II LRs of *ckx2* loss-of-function mutant plants showed a significant reduction of meristem size at the upper LR flank compare to wild-type Col-0 (Figure 7F). Similarly, BAP treatment resulted in the development of shorter meristems in stage II LRs of Col-0 wild-type (Figure S7F). These data suggest that CK signaling also negatively regulates meristem activity in emerged LRs.

We next used cell division marker *pCycB1;1::GUS*, to assess the spatial impact of CK on cell proliferation. BAP and INCYDE treatment reduced the abundance of *CycB1;1::GUS* at the upper flank of stage II LRs (Figure S7G-H). These data suggest that CK signaling does not only restrict cellular elongation, but also reduces cell proliferation in emerged LRs.

Notably, *CDKB1;1* and other cell cycle promoting genes are down-regulated in the *crf1,3,5,6* quadruple mutant (Raines et al., 2015). Hence, we assumed that CRF-dependent control of the cell cycle may contribute to the CK-mediated establishment of GSA in emerged LRs. To block cell cycle progression, we used the dominant negative (DN) allele of CDKB1;1 and the *cdkb1;1 cdkb1;2* double mutant (Figure 7G), as well as the cell cycle inhibitor Roscovitine (Figure 7H). Both genetic and pharmacological interference with the cell cycle strongly interfered with the LR bending (Figure 7G-H). In contrast to LRs, the gravity response kinetics in primary roots of CDKB1;1^DN^ as well as *cdkb1;1 cdkb1;2* were similar to wild-type behavior (Figure S7I). This suggests that not only cellular elongation (Rosquete et al., 2013), but also cell proliferation in stage II LRs is a particular determinant of directional LR growth.

Overall, these data suggest that CK modulates both differential cell elongation and cell proliferation to interfere with growth at the upper flank of LR, ultimately regulating angular LR growth and radial expansion of the root system.

## Discussion

Because root systems are hidden beneath the soil, the study and directed improvement of root architectural traits in crop breeding programs have been delayed. There is growing interest to alleviate the harmful effects of drought stress by modulating the primary GSA of LRs (Uga et al., 2013). Despite the apparent importance of the root system depth, the molecular mechanisms regulating the direction of LR growth are poorly understood. Thus, understanding the molecular mechanisms establishing the primary angular growth in LRs could guide future engineering of plants to suit certain habitats. Anticipating that natural variation could provide valuable insights on how to sustainably engineer root systems, we focused on the primary growth direction of lateral roots in natural *Arabidopsis* accessions.

We reveal that the primary growth direction of lateral roots varies substantially within a population of natural *Arabidopsis* accessions. Primary LR angles of hypo- or hyper-responsive accessions followed a similar trend regardless of whether they were grown in soil, two-dimensional or three-dimensional in-vitro systems. We thus conclude that this approach is suitable to assess the genetic control of angular LR growth. Using a GWAS approach, we show that angular growth of lateral roots is controlled by CKX2- dependent metabolism of the phytohormone cytokinin. CKX2 contains a SP to enter the secretory pathway, which could be crucial for its impact on CK perception. However, the precise site of CK receptor activity (plasma membrane and/or endoplasmic reticulum) is still under debate (Romanov et al., 2018). We conclude that variation in an amino acid substitution after the predicted cleavage site impacts on SP processing of CKX2, which consequently obstructs CKX2 activity *in planta*. Our data suggest that the lack of SP processing abolishes the secretion and enzymatic activity of CKX2, thereby contributing to CK-dependent GSA trait variation in natural Arabidopsis accessions.

Nordic accessions preferentially express an inactive *CKX2* variant, which prompted us to investigate whether environmental cues further define the root system in a CK-dependent manner. We revealed that hypoxic conditions induce more horizontal LR growth through CK signaling. The increased frequency of an inactive *CKX2* allele in Nordic accessions suggests that the allele may have been selected for in these populations, promoting more horizontal root growth. It is an intriguing possibility that more horizontal, near surface roots may rectify gas exchange under hypoxia conditions, potentially alleviating the harmful effects of hypoxic stress in these Nordic, snow covered habitats.

Our analysis suggests that primary and secondary roots have distinct responses to CK. While CK signaling abolishes PIN-dependent transport in main roots (Marhavý et al., 2011), we showed that CKX2-dependent interference with endogenous CK levels does not affect PIN3 and auxin signaling in emerged LRs. Moreover, CK signaling is asymmetric in emerged lateral, but not primary roots, proposing a unique role of cytokinin in regulating asymmetric growth responses in LRs. Also abscisic acid signaling displays distinct activities in main and lateral root organs, presumably allowing distinct organ growth rates in response to environmental stresses (Ding and De Smet, 2013). Thus, we propose that hormone signaling might be generally co-opted in primary and secondary roots to facilitate different growth responses to the environment.

The increase and decrease of cytokinin levels have been shown to slightly accelerate the rate of gravitropic bending in primary root, but the developmental importance of this effect remains uncertain (Pernisova et al., 2016). In contrast, we show here that CK signaling plays a developmental role in establishing the primary GSA of LRs. Moreover, an increase and decrease of CK signaling correlate with reduced and enhanced down-ward bending of LRs, respectively. Mechanistically, we showed that CK signaling interferes with cellular elongation and proliferation in emerged LR to reduce LR organ bending towards gravity. These stage II LRs undergo a *de novo* formation of the elongation zone (Rosquete et al., 2013). During this developmental time window, the CK-dependent reduction in cell proliferation could have hence an immediate influence on the number of elongating cells. Such an impact could further compromise angular LR growth, because our computational model predicted that an asymmetric reduction in cell number (at the upper root flank) would induce mechanical constraints, additionally limiting organ bending (Figure S7C-D). However, such detailed mechanical constraint measurements in LRs await experimental validation.

We illustrate that CKX2 contributes to the rate of asymmetric CK signaling, but *CKX2* expression did not show a pronounced asymmetry. Similarly, the CK response factors CRF2 and CRF3 are not asymmetrically expressed in emerged lateral roots. Thus, the molecular mechanism by which asymmetric CK signalling across the LR tip is established remains to be elucidated. Our work proposes that an auxin transport mechanism promotes the asymmetry of CK signaling. Accordingly, auxin could generate an anti-gravitropic signal to interfere with its own gravitropic impact in LRs. Unlike auxin, the mechanisms of intercellular cytokinin transport are poorly characterized (Kang et al., 2017). One intriguing possibility is however that the asymmetric auxin signal could favor CK relocation towards the upper side of LRs, inducing differential CK signalling and growth repression on this side. However, it is also possible that differential CK signaling occurs at the level of signal integration and might be independent of differential distribution of CK. Future work will examine these possibilities to uncover the mechanism by which CKX2 is linked to differential cytokinin activity across a stage II LR.

In conclusion, our genetic screen uncovered that directional LR growth depends on opposing gravitropic and anti-gravitropic phytohormonal cues (Figure S7J). We conclude that CK signaling reduces growth at the upper organ side, which counteracts the gravity induced, auxin-dependent reduction in cell expansion at the lower root flank. In this way, a CK-dependent mechanism allows the root system to override the gravitropic response and radially explore its surroundings. Genetic interference with CK signaling cannot only be used to define the primary growth direction of LRs, but moreover may refract certain environmental input to root architecture. Overall, these results propose that directed interference with CK responses in LRs could be used to engineer root system depth to better suit certain habitats.

## Acknowledgments

We are grateful to Bruno Müller, Thomas Schmülling, Magnus Nordborg, Wolfgang Busch, Ben Scheres, Jiri Friml, Dirk Inze, Tomas Werner, Marketa Pernisova, Eva Benkova, Joseph Kieber and Lieven De Veylder for sharing published material; Marget Sauter, Ilka Reichardt-Gomez, Ümit Seren and Envel Kerdaffrec for helpful discussions; Jit Thacker for help with preparing the manuscript; Hana Martínková for help with phytohormone analyses; and the BOKU-VIBT Imaging Centre for access and expertise. This work was supported by the Austrian Academy of Sciences (ÖAW) (DOC fellowship to K.D.), Fulbright-Austria Marshall Plan student grant (to E.S.), Vienna Research Group (VRG) program of the Vienna Science and Technology Fund (WWTF) (to J.K-V.), the Austrian Science Fund (FWF) (P29754) (to J.K-V.), the European Research Council (ERC) (Starting Grant 639478-AuxinER) (to J.K-V.), and work was funded by the Ministry of Education, Youth and Sports of the Czech Republic (National Program for Sustainability I, grant no. LO1204) (to O.N.), and Programa de Atracción de Talento 2017 (Comunidad de Madrid, 2017-T1/BIO-5654 to K.W.).

## Author contributions

S.W. performed most experiments. M.R.R. initiated the project. M.S., E.S. and K.D. performed confocal microscopy. H.L., T.L.R. and J.R.D. contributed GLO-Roots data. I.P. and O.N. conducted quantification of endogenous cytokinins. S.M. and R.S. performed hypoxia experiments K.W. designed and described the dynamic computer model simulation. J.K.-V. devised and coordinated the project. S.W. and J.K.-V. wrote the manuscript. All authors saw and commented on the manuscript.

## Material and methods

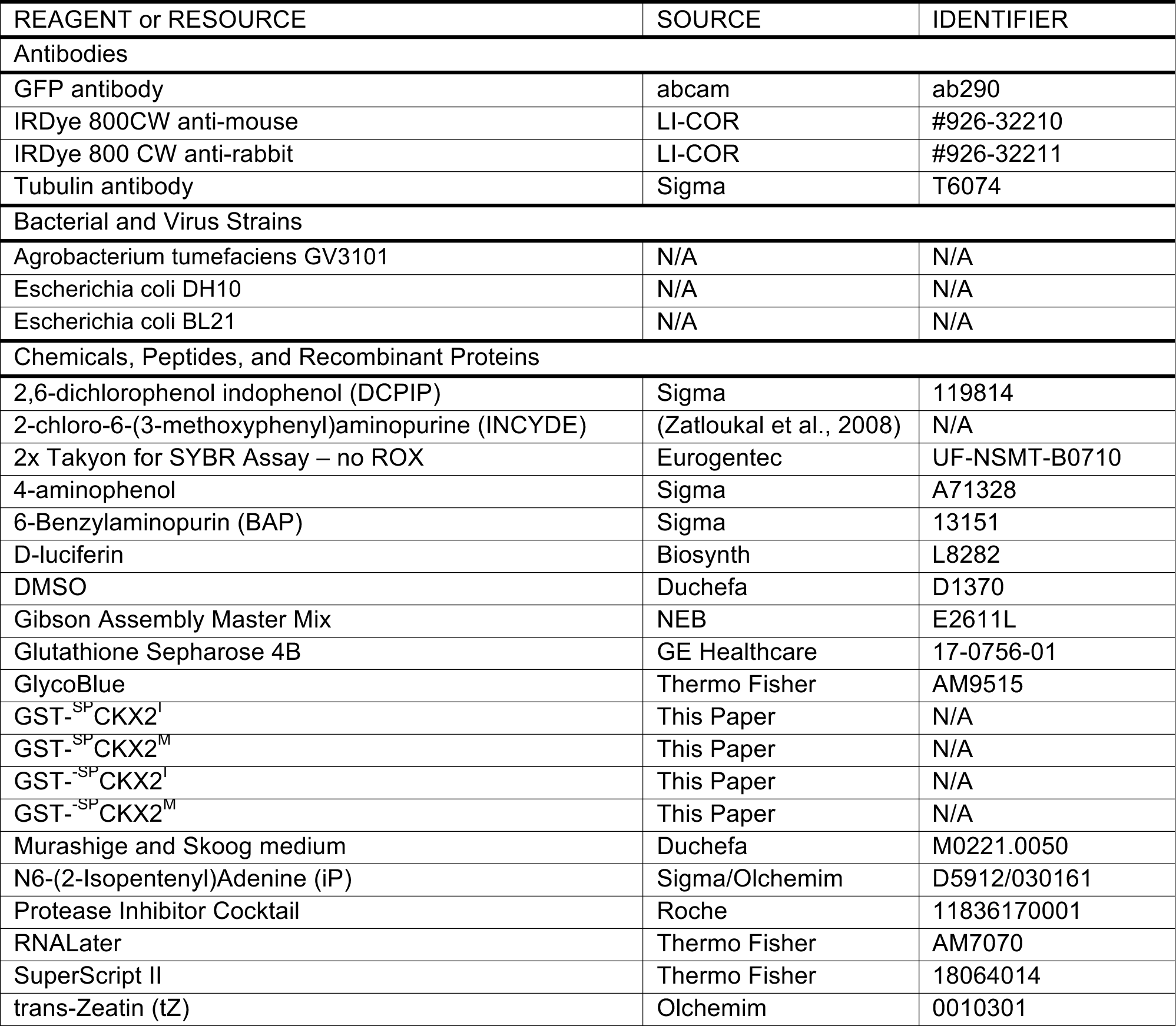
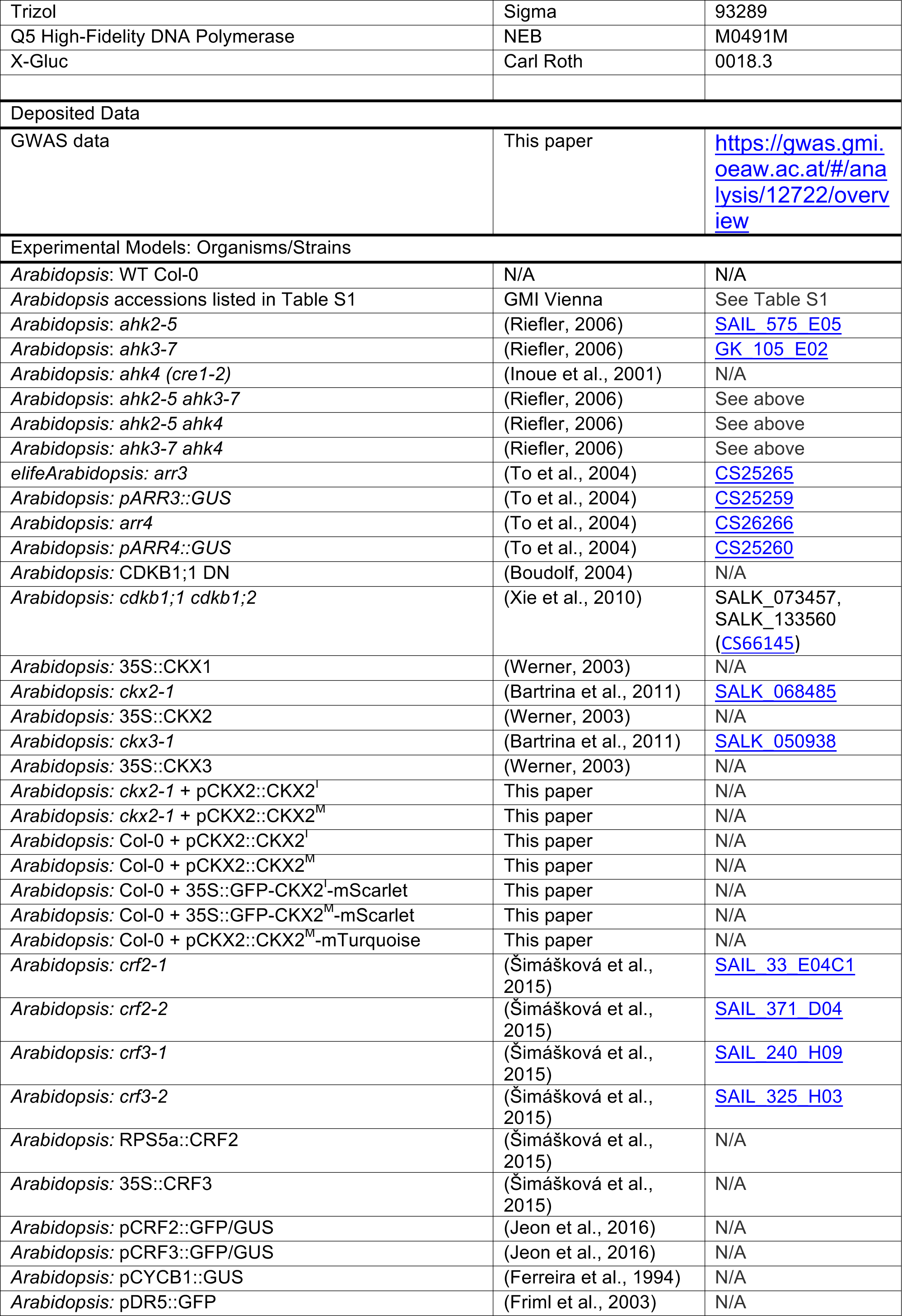
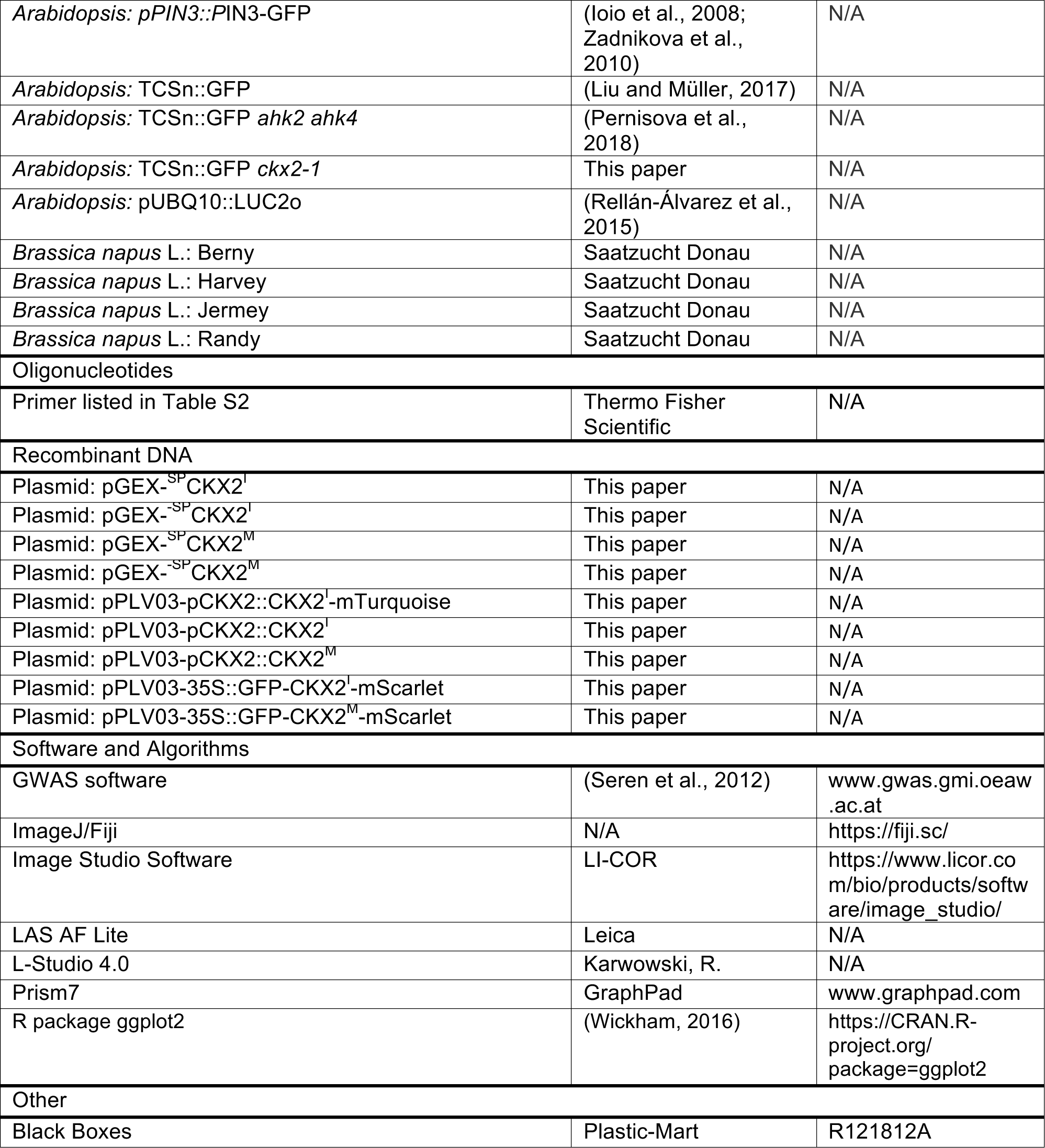
KEY RESOURCES TABLE.

### Plant material and growth conditions

Seeds of *Arabidopsis thaliana* accessions were kindly provided by Magnus Nordborg and Wolfgang Busch. A list of all lines used can be found in Supporting Information, Table S1. Seeds of *Brassica napus* L. were kindly provided by Saatzucht Donau. Seeds were surface sterilized, stratified at 4°C for 2 days in the dark. Seedlings were grown vertically on half Murashige and Skoog medium (1/2 MS salts (Duchefa), pH 5.9, 1% sucrose, and 0.8% agar). Plants were grown under long-day (16 h light/8 h dark) conditions at 20–22°C.

### Chemicals and Treatments

6-Benzylaminopurin (BAP) (Sigma), trans-Zeatin (tZ) (OlChemim), N^6^-(2-Isopentenyl)Adenine (iP) (OlChemim) were all dissolved in DMSO (Duchefa). 2-chloro-6-(3-methoxyphenyl)aminopurine (INCYDE) was synthesized in by the Laboratory of Growth Regulators, Palacký University & Institute of Experimental Botany AS CR (Olomouc, Czech Republic) as previously described (Zatloukal et al., 2008) and dissolved in DMSO. Treatments with BAP, tZ, iP and INCYDE were all performed on 7 day old seedlings (transferred to supplemented media).

GUS stainings were performed after 24h and initial GSA measurements 7 days after transfer.

### Genome-wide association studies (GWAS)

To identify the genetic basis of the for the GSA of LRs we carried out a GWAS using an accelerated mixed model (AMM) (Seren et al., 2012). The GWAS results can be viewed interactively online: https://gwas.gmi.oeaw.ac.at/#/analysis/12722/overview.

### GLO-Roots

#### Rhizotrons

The basic rhizotron design was as described in (Rellán-Álvarez et al., 2015). To adapt the rhizotrons for use in an automated rhizotron handling system (designed by Modular Science, San Francisco), several modifications were implemented. The top edge of each rhizotron sheet was beveled using a belt sander to facilitate automated watering. Two 1/16” thin aluminium hooks used for automatic handling of the rhizotron were attached on each side of the rhizotron. To reduce light exposure of the root system during growth, a 1/8” thin black acrylic rhizotron top shield was installed.

#### Boxes and holders

Black 12” W × 18” L × 12” H boxes (Plastic-Mart) were used to grow plants in 12 rhizotrons at a time. The arrangement of 1/8”-thin black acrylic sheets of different shapes and sizes formed 12 light-proof chambers to make sure that the roots of every rhizotron were shielded from light even when one rhizotron was removed for imaging. Rhizotron preparation

Rhizotron preparation was as described in (Rellán-Álvarez et al., 2015) with slight modifications required by the new rhizotron design.

#### Plant growth in rhizotrons

Two transfer pipettes (each ∼2ml) of quick-releasing fertilizer (Peter’s 20-20-20) were added to each rhizotron after assembly. Assembled rhizotrons were placed into a box with water and allowed to absorb water overnight from the bottom side. Seeds containing the pUBQ10:LUC2o transgene were stratified for 2 days at 4°C in distilled water and 3 seeds were sown in the center of each rhizotron. Each rhizotron was equipped with a unique barcode. All rhizotrons were sprayed down with water and sealed with a transparent lid and packing tape. Plants were grown at 22/18°C (day/night) under long-day conditions (16h light, 8h dark) using LED lights (Vayola, C-Series, N12 spectrum) with a light intensity of about 130 µmol m^-2^ s^-1^. After 2 days, the transparent lid was unsealed, rhizotrons were watered with 2 transferring pipettes of water, and the lid left loose for an additional day. After removing the lid, rhizotrons were watered twice per day with 2 transferring pipettes of water each time until 9 days after sowing. Plant imaging:

20 days after sowing, the automated rhizotron handling system (designed by Modular Science, San Francisco) added 50ml of 300µM D-luciferin (Biosynth) at the top of each rhizotron and loaded the rhizotron into a fixed stage that was controlled by a Lambda 10-3 optical filter changer (Sutter Instruments, Novato, CA) in the GLO1 imaging system (Rellán-Álvarez et al., 2015). 5-min exposures were taken per rhizotron side. A shoot image was taken right after the four root images using an ids UI-359xLE-C camera with a Fujinon C-Mount 8-80mm Varifocal lens that was installed in GLO1. Three LED strips on each side of the camera were switched on before a shoot image was taken.

#### Image preparation

Image preparation was similar to in (Rellán-Álvarez et al., 2015): four individual root images were collected: top front, bottom front, top back, and bottom back. Using an automated ImageJ macro, a composite image was generated as follows: (1) images were rotated and translated to control for small misalignments between the two cameras; (2) the top and bottom images of each side were merged; (3) the back image was flipped horizontally; (4) the front and back images were combined using the maximum values. The final images produced were 16-bit in depth and 4096 × 2048 pixels. The scale of the images was 138.6 pixels per cm.

### Hypoxia Treatment

All following treatments were performed in air-tight glass desiccators in which seedlings grown on vertical agar plates were carefully placed with the lids removed. Seedlings were exposed to a hypoxia treatment by flushing the desiccators with humidified 100% N_2_ gas (2l/min) for 4 hours (13.00-17.00h) in the dark to limit photosynthesis derived oxygen production. For the controls, desiccators were flushed with humidified air. Flow rates were controlled by mass flow controllers (MASS-VIEW, Bronkhorst). At the end of the hypoxia treatment, plates were carefully removed from the desiccators, closed, and transferred back to the climate chamber. The plates remained in the climate chambers under control growth conditions for 5 days after the treatment after which they were scanned using an EPSON Scanner V300.

### DNA constructs

The promoter region and full-length CKX2^I^ or coding DNA sequence (CDS) were amplified by PCR (Table S2) from genomic DNA or cDNA using Q5 High-Fidelity DNA Polymerase (NEB) and cloned either alone or under of the 35S promoter together with GFP and mScarlet-i into pPLV03 or pGEX5x3 using Gibson Assembly (NEB). Subsequently, this plasmid were used for *in vitro* mutagenesis (Table S2) to obtain CKX2^M^. The resulting constructs were transformed into Col-0 and *ckx2-1* plants using the floral dipping method (Clough and Bent, 1998) or for transient transformation in tobacco plants.

### Expression, Purification and Activity Measurement of Recombinant Proteins

Recombinant proteins were expressed as GST fusion proteins and in *Escherichia coli* BL21 codon plus strain. Proteins were purified using the Sepharose beads affinity method (Glutathione Sepharose 4B; GE Healthcare).

The activity was measured using a modified end-point method previously described (Frébort et al., 2002). For activity screening, the samples were incubated in a reaction mixture (total volume of 600 µl) that consisted of 200 mM McIlvaine buffer (100 mM citric acid and 200 mM Na_2_HPO_4_) pH 7.4, 500 μM 2,6-dichlorophenol indophenol (DCPIP; Sigma) as electron acceptor and different concentrations of N^6^-(2-isopentenyl)adenine (iP; Sigma) as substrate. The volume of the enzyme sample used for the assay was adjusted based on the enzyme activity. The incubation time at 37 °C was 1h. The enzymatic reaction was stopped after incubation by adding 300 µl of 40% trichloroacetic acid (TCA), then 200 µl 2% 4-aminophenol (Sigma) (in 6% TCA) was added and the sample was centrifuged at 13,200 rpm for 5 min to remove protein precipitate. 200 µl supernatant was used to measure the absorption spectrum from 352 nm to 500 nm to determine the concentration of produced Schiff base with ε352=15.2 mM^−1^cm^−1^ using a plate reader.

### Microscopy

Confocal microscopy was performed using a Leica SP5 (Leica). Fluorescence signals for GFP (excitation 488 nm, emission peak 509 nm), mScarlet-i (excitation 561 nm, emission peak 607 nm) and propidium iodide (PI) staining (excitation 569 nm, emission peak 593 nm) were detected with a 40x or 63x (water immersion) objective. Image processing was performed using LAS AF lite software (Leica). Graphpad Prism software was used to evaluate the statistical significance of the differences observed between control and treated groups (One-way ANOVA).

### Gravitropic Set-Point Angle Measurements

Plates with 14-day-old seedlings were scanned and the initial gravitropic set-point angle (iGSA) of individual LRs was measured with reference to the gravity vector (Rosquete et al., 2013) using Image J software. Individual GSA values were then sorted into 8 categories: 0 °-30 °, 31 °-50 °, 51 °-70 °, 71 °-90 °, 91 °-110 °, 111 °-180 °. Percentages of incidence were calculated for each category and graphs of GSA distribution were generated. The test of Kolmogorov-Smirnov (KS-test) was used online (http://www.physics.csbsju.edu/stats/KS-test.n.plot_form.html) to statistically evaluate the GSA data sets generated from mutants and treated seedlings in comparison to wild type and untreated controls, respectively.

### Histochemical GUS Staining

GUS histochemical staining of acetone-fixed 7-day-old seedlings containing pCycB1::GUS fusion constructs followed a previously described method (Crone et al., 2001) using x-Gluc (Carl Roth) as substrate. Examination of stained seedlings and image acquisition were performed with a light microscope (Zeiss Observer D1) equipped with a DFC 300 FX camera (Zeiss). Graphpad Prism software was used to evaluate the statistical significance of the differences observed between control and treated groups (One-way ANOVA).

### Transient Transformation, Protein Extraction and Immunoblot Analysis

The *Agrobacterium tumefaciens* strain GV3101 was transformed with the respective construct and grown for 2 d at 28°C in 5 ml Luria-Bertani (LB). The preculture was used to inoculate 25 ml LB and incubated for 4 h at 28°C. Cells were pelleted and resuspended in 30 ml LB supplemented with 100 μM acetosyringone. After 2 h, cells were resuspended in 30 ml of 5% sucrose and infiltrated in tobacco (*Nicotiana tabacum*) leaves. Subcellular localization was examined 3 d after transformation by confocal laser scanning microscopy (see above) or leaves were ground to fine powder in liquid nitrogen and solubilized with extraction buffer (25 mM TRIS, pH 7.5, 10 mM MgCl_2_, 15 mM EGTA, 75 mM NaCl, 1 mM DTT, 0.1% Tween20, with freshly added proteinase inhibitor cocktail (Roche)). Protein concentration was assessed using the Bradford method. Membranes were probed with a 1:1,000 dilution of GFP antibody (abcam) or 1:20,000 of tubulin antibody (Sigma). Goat IRDye 800CW anti-mouse (LI-COR) or goat IRDye 800 CW anti-rabbit (LI-COR) was used (1:20,000) as secondary. The signals were detected and quantified using the Odyssey Imagine System (LI-COR).

### RNA Extraction, cDNA Synthesis and Quantitative PCR

RNA was extraction was describe previously (Hofmann et al., 2019). In brief: A pool of 10 lateral roots or root tips were collected in 30 µl of 100% RNAlater (Thermo Fisher) and 500 µl of TRIzol (Sigma) was added followed by brief vortexing (2 × for 2 s each) and incubating at 60°C for 30 min. 100 µl of chloroform was added, and then, samples were vortexed briefly (2 × for two seconds each) and incubated at room temperature for three minutes. After centrifugation at 12,000*g* for 15 min at 4°C, the aqueous phase was transferred to a new LoBind tube. To precipitate the RNA, an equal volume of isopropanol and 1.5 µl of GlycoBlue (Thermo Fisher) was added followed by a − 20°C incubation for 15–18 h and centrifugation at > 20,000 g for 30 min at 4°C. After removal of the supernatant, the pellet was washed by adding 500 µl of 75% ethanol, vortexing briefly and then centrifuged at > 20,000 g for 15 min at 4°C. The 75% ethanol wash step was repeated 1 ×. As much ethanol as possible was removed followed by the drying of the pellet by letting the Eppendorf tube sit on ice with lid open for 10 min. Precipitated RNA was then resuspended with 5–12 µl of nuclease-free water, stored at − 80°C. cDNA synthesis was performed using SuperScript II (Thermo Fisher) and qPCR using 2x Takyon for SYBR Assay – no ROX (Eurogentec) following the manufactures instructions on a CFX96 Touch Real-Time PCR Detection System (Bio-Rad). Expression values were normalized to the expression of ubiquitin 5 (UBQ5) and translation initiation factor EIF4A.

### Cytokinin Measurements

Quantification of cytokinin metabolites was performed according to the method described by (Svačinová et al., 2012), including modifications described in (Antoniadi et al., 2015). Briefly, samples (20 mg FW) were extracted in 1 ml of modified Bieleski buffer (Hoyerová et al., 2006) together with a cocktail of stable isotope-labeled internal standards used as a reference (0.25 pmol of CK bases, ribosides, N-glucosides, and 0.5 pmol of CK O-glucosides, nucleotides per sample added). The extracts were purified using the Oasis MCX column (30 mg/1 ml, Waters) and cytokinin levels were determined using the LC-MS/MS system consisting of an ACQUITY UPLC System and a Xevo TQ-S triple quadrupole mass spectrometer (Waters). Results are presented as the average of five biological replicates ± standard deviation in pmol/g FW. Statistical examinations were made between Col-0 wild type and *ckx2-1* roots using two-way ANOVA analysis.

### Description of the computer model of LR

For the sake of simplicity, our model is composed of a rectangular grid in which each box represents a single cell. Cell walls are modelled as a linear elastic spring (connecting 2 adjacent vertices) that can expand and contract in order to minimize forces acting on each spring. The magnitude of force exerted by this spring is 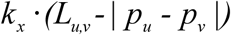 and is positive for spring compression. The *k_x_* characterizes the stiffness of the spring and was set to 0.9 in all simulations. This force is in the direction of the spring 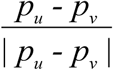. *p_u_* is the position of vertex *u,* and *p_v_* is the position of neighbour vertex *v.* The total force exerted on vertex *u* located at position *p_u_* by all such springs can be written as:

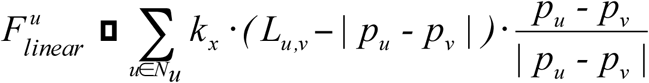

where *N_u_* is the set of vertices adjacent to vertex *u*. The norm symbol indicates the Euclidean distance between the points.

In addition to the forces acting on a vertex due to springs, a force due to the turgor pressure inside the cell (*p_const_* = 0.005) acts in the direction normal to each wall (***n***):

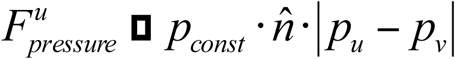

Combining the individual force components, the total force acting on a vertex *u* is the sum of forces acting on each cell wall and internal pressure inside the cells.

In accordance with Newton’s Second Law of motion we calculated the velocity (*Vel_u_*) and position (*p_u_*) of vertex *u* over time for point mass *m*_u_ = 1 with the following formulas:

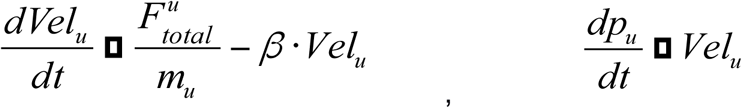

where *β=0.2 is* a damping constant.

The LR root model is spatially divided into two zones (root tip and elongation zones) (Figure S9) along the x-axis based on the threshold parameter that defines distance between individual cell centers and right-most cell centers. This parameter controls the length of the elongation zone or simply the number of elongating cells in our simulations (Figure S8C-D). Along the y-axis, cells elongate at different rates based on the smooth polynomial interpolation between the minimum measured elongation rate at the bottom part of the LR (5 µm/h) to the maximum elongation rate at the top part of the LR (15 µm/h) (Rosquete et al., 2013). Cell elongation is simulated by expanding the resting length of linear springs.

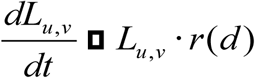

Where *r(d)* is an interpolated growth rate and *d* is the relative distance from the bottom part of the LR such that *r_min_(0)* = 5 µm/h and *r_max_(1)* = 15 µm/h.

The geometry of the model was created using a version of the VV simulator (Smith et al., 2003; 2006) embedded in the modeling software L-studio (Karwowski) (http://algorithmicbotany.org/lstudio). Cell mechanics (mass-spring system) was simulated using the forward Euler method. All simulations were stopped after 8 hours of elapsed growth as observed experimentally. Screenshots from model simulations are shown in Figure S9A.

## Supplemental Information

**Table S1.**
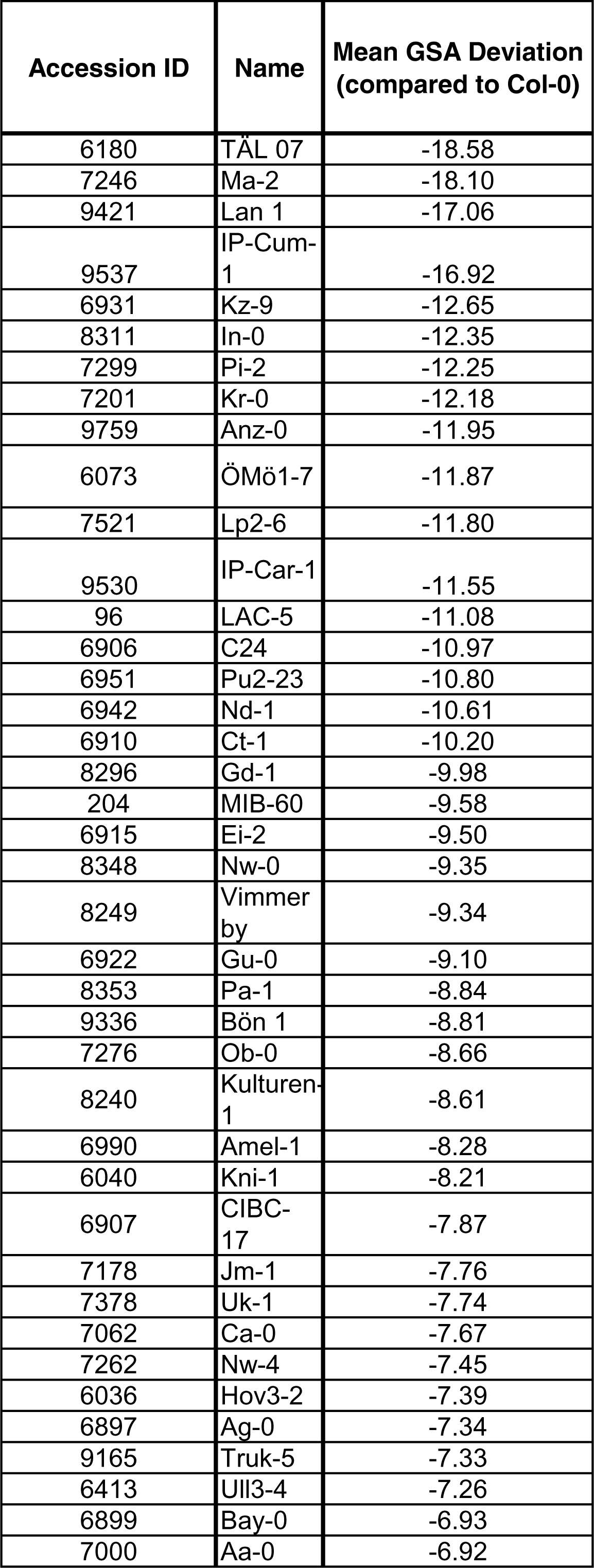
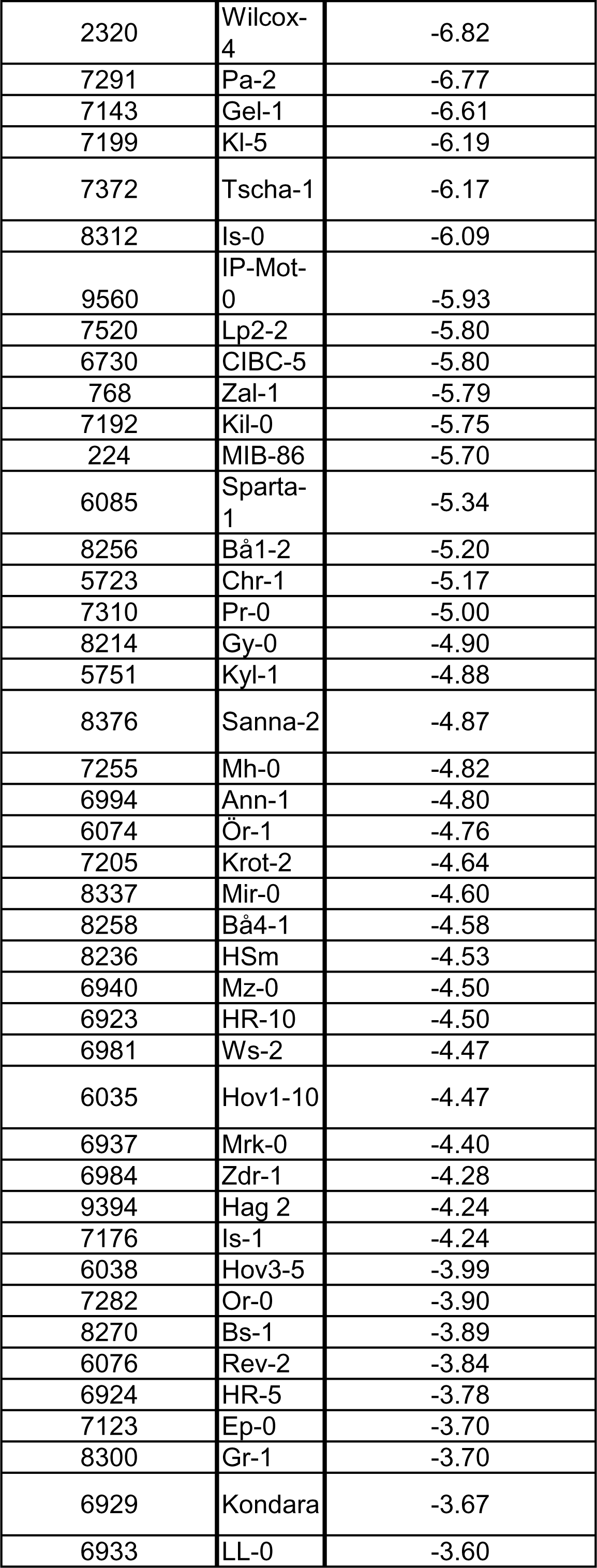
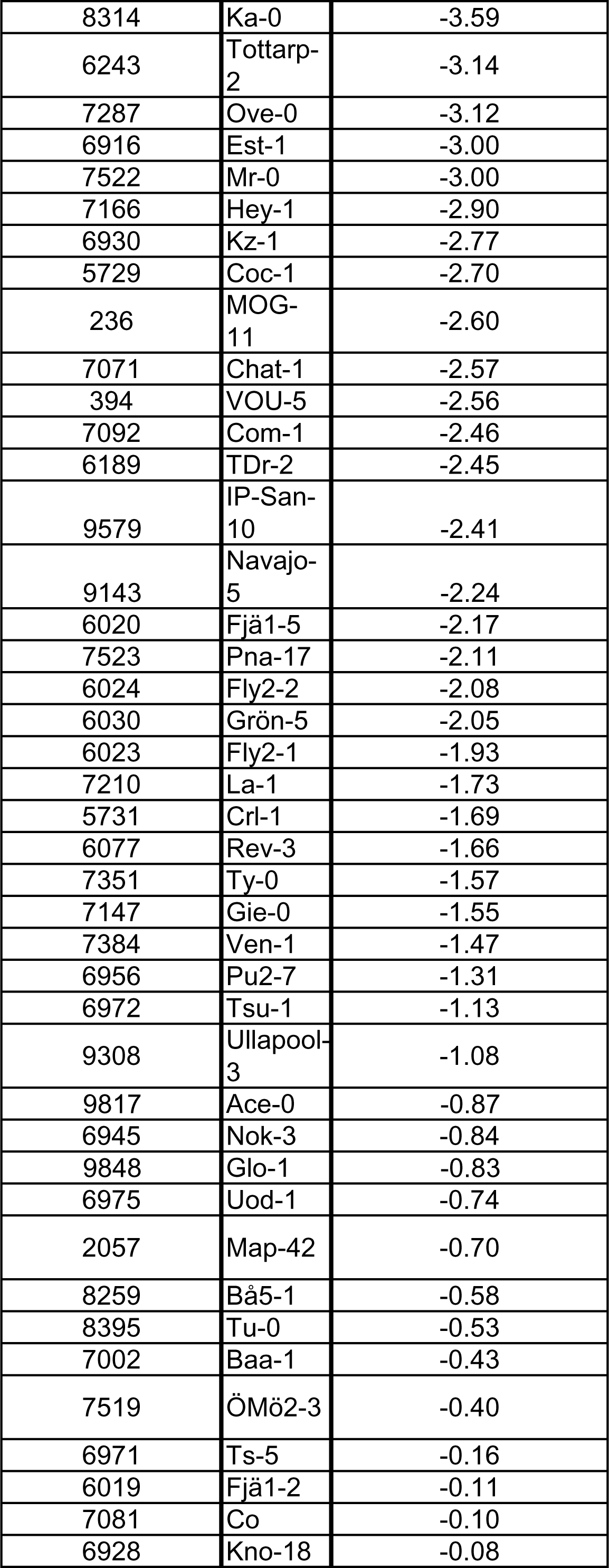
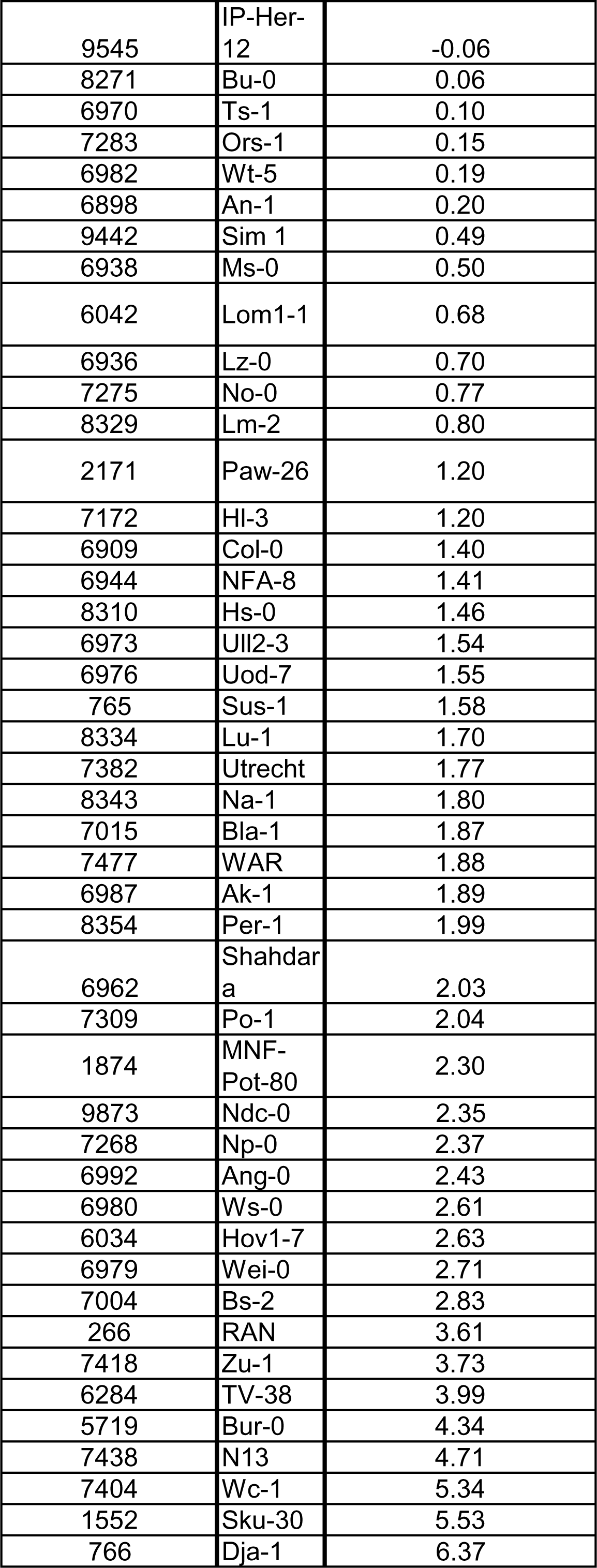
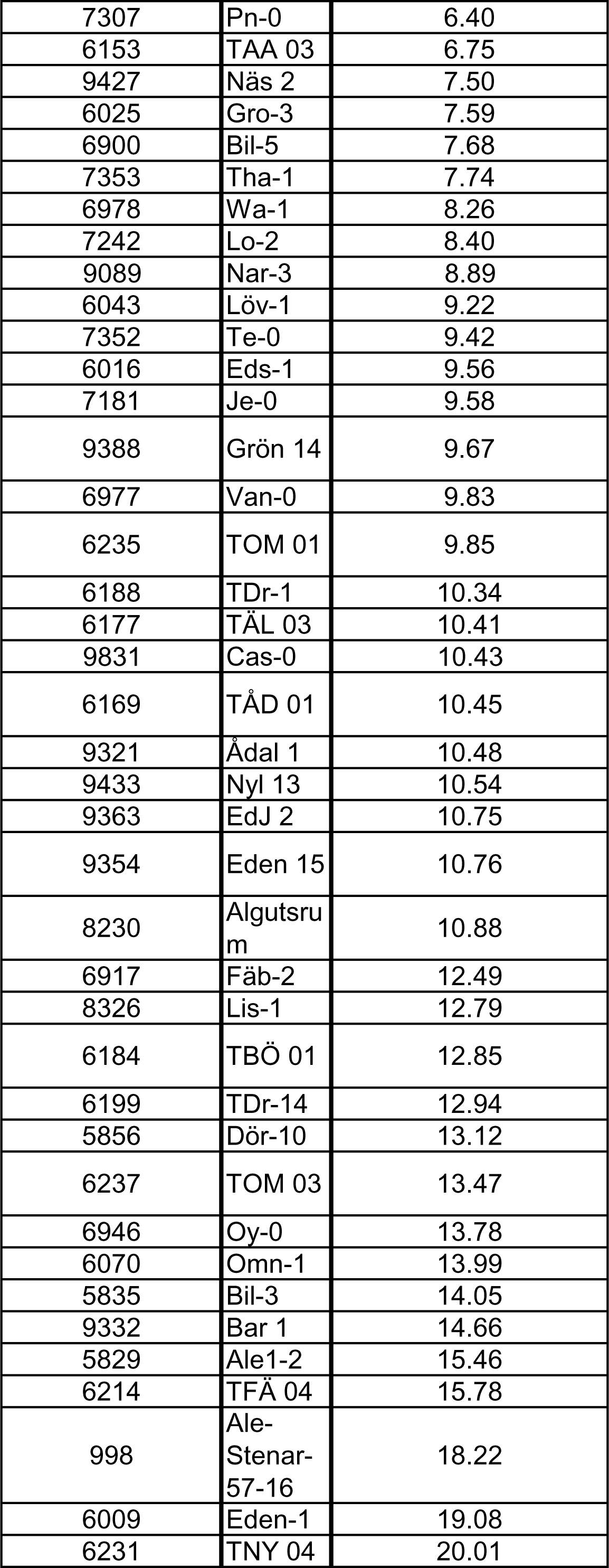
Accessions with their mean GSA distribution (compared to Col-0) used in this study.

**Figure S7. Cytokinin-dependent interference with cell division rates defines angular growth of lateral roots.**
| Accession ID | Name | Mean GSA Deviation<br>(compared to Col-0) |
| --- | --- | --- |
| 6180 | TÄL 07 | -18.58 |
| 7246 | Ma-2 | -18.10 |
| 9421 | Lan 1 | -17.06 |
| 9537 | IP-Cum-1 | -16.92 |
| 6931 | Kz-9 | -12.65 |
| 8311 | In-0 | -12.35 |
| 7299 | Pi-2 | -12.25 |
| 7201 | Kr-0 | -12.18 |
| 9759 | Anz-0 | -11.95 |
| 6073 | ÖMö1-7 | -11.87 |
| 7521 | Lp2-6 | -11.80 |
| 9530 | IP-Car-1 | -11.55 |
| 96 | LAC-5 | -11.08 |
| 6906 | C24 | -10.97 |
| 6951 | Pu2-23 | -10.80 |
| 6942 | Nd-1 | -10.61 |
| 6910 | Ct-1 | -10.20 |
| 8296 | Gd-1 | -9.98 |
| 204 | MIB-60 | -9.58 |
| 6915 | Ei-2 | -9.50 |
| 8348 | Nw-0 | -9.35 |
| 8249 | Vimmer<br>by | -9.34 |
| 6922 | Gu-0 | -9.10 |
| 8353 | Pa-1 | -8.84 |
| 9336 | Bön 1 | -8.81 |
| 7276 | Ob-0 | -8.66 |
| 8240 | Kulturen-1 | -8.61 |
| 6990 | Amel-1 | -8.28 |
| 6040 | Kni-1 | -8.21 |
| 6907 | CIBC-17 | -7.87 |
| 7178 | Jm-1 | -7.76 |
| 7378 | Uk-1 | -7.74 |
| 7062 | Ca-0 | -7.67 |
| 7262 | Nw-4 | -7.45 |
| 6036 | Hov3-2 | -7.39 |
| 6897 | Ag-0 | -7.34 |
| 9165 | Truk-5 | -7.33 |
| 6413 | UII3-4 | -7.26 |
| 6899 | Bay-0 | -6.93 |
| 7000 | Aa-0 | -6.92 |
| 2320 | Wilcox-4 | -6.82 |
| 7291 | Pa-2 | -6.77 |
| 7143 | Gel-1 | -6.61 |
| 7199 | KI-5 | -6.19 |
| 7372 | Tscha-1 | -6.17 |
| 8312 | Is-0 | -6.09 |
| 9560 | IP-Mot-0 | -5.93 |
| 7520 | Lp2-2 | -5.80 |
| 6730 | CIBC-5 | -5.80 |
| 768 | Zal-1 | -5.79 |
| 7192 | Kil-0 | -5.75 |
| 224 | MIB-86 | -5.70 |
| 6085 | Sparta-1 | -5.34 |
| 8256 | Bå1-2 | -5.20 |
| 5723 | Chr-1 | -5.17 |
| 7310 | Pr-0 | -5.00 |
| 8214 | Gy-0 | -4.90 |
| 5751 | Kyl-1 | -4.88 |
| 8376 | Sanna-2 | -4.87 |
| 7255 | Mh-0 | -4.82 |
| 6994 | Ann-1 | -4.80 |
| 6074 | Ör-1 | -4.76 |
| 7205 | Krot-2 | -4.64 |
| 8337 | Mir-0 | -4.60 |
| 8258 | Bå4-1 | -4.58 |
| 8236 | HSm | -4.53 |
| 6940 | Mz-0 | -4.50 |
| 6923 | HR-10 | -4.50 |
| 6981 | Ws-2 | -4.47 |
| 6035 | Hov1-10 | -4.47 |
| 6937 | Mrk-0 | -4.40 |
| 6984 | Zdr-1 | -4.28 |
| 9394 | Hag 2 | -4.24 |
| 7176 | Is-1 | -4.24 |
| 6038 | Hov3-5 | -3.99 |
| 7282 | Or-0 | -3.90 |
| 8270 | Bs-1 | -3.89 |
| 6076 | Rev-2 | -3.84 |
| 6924 | HR-5 | -3.78 |
| 7123 | Ep-0 | -3.70 |
| 8300 | Gr-1 | -3.70 |
| 6929 | Kondara | -3.67 |
| 6933 | LL-0 | -3.60 |
| 8314 | Ka-0 | -3.59 |
| 6243 | Tottarp-2 | -3.14 |
| 7287 | Ove-0 | -3.12 |
| 6916 | Est-1 | -3.00 |
| 7522 | Mr-0 | -3.00 |
| 7166 | Hey-1 | -2.90 |
| 6930 | Kz-1 | -2.77 |
| 5729 | Coc-1 | -2.70 |
| 236 | MOG-11 | -2.60 |
| 7071 | Chat-1 | -2.57 |
| 394 | VOU-5 | -2.56 |
| 7092 | Com-1 | -2.46 |
| 6189 | TDr-2 | -2.45 |
| 9579 | IP-San-10 | -2.41 |
| 9143 | Navajo-5 | -2.24 |
| 6020 | Fjä1-5 | -2.17 |
| 7523 | Pna-17 | -2.11 |
| 6024 | Fly2-2 | -2.08 |
| 6030 | Grön-5 | -2.05 |
| 6023 | Fly2-1 | -1.93 |
| 7210 | La-1 | -1.73 |
| 5731 | CrI-1 | -1.69 |
| 6077 | Rev-3 | -1.66 |
| 7351 | Ty-0 | -1.57 |
| 7147 | Gie-0 | -1.55 |
| 7384 | Ven-1 | -1.47 |
| 6956 | Pu2-7 | -1.31 |
| 6972 | Tsu-1 | -1.13 |
| 9308 | Ullapool-3 | -1.08 |
| 9817 | Ace-0 | -0.87 |
| 6945 | Nok-3 | -0.84 |
| 9848 | Glo-1 | -0.83 |
| 6975 | Uod-1 | -0.74 |
| 2057 | Map-42 | -0.70 |
| 8259 | Bå5-1 | -0.58 |
| 8395 | Tu-0 | -0.53 |
| 7002 | Baa-1 | -0.43 |
| 7519 | ÖMö2-3 | -0.40 |
| 6971 | Ts-5 | -0.16 |
| 6019 | Fjä1-2 | -0.11 |
| 7081 | Co | -0.10 |
| 6928 | Kno-18 | -0.08 |
| 9545 | IP-Her-12 | -0.06 |
| 8271 | Bu-0 | 0.06 |
| 6970 | Ts-1 | 0.10 |
| 7283 | Ors-1 | 0.15 |
| 6982 | Wt-5 | 0.19 |
| 6898 | An-1 | 0.20 |
| 9442 | Sim 1 | 0.49 |
| 6938 | Ms-0 | 0.50 |
| 6042 | Lom1-1 | 0.68 |
| 6936 | Lz-0 | 0.70 |
| 7275 | No-0 | 0.77 |
| 8329 | Lm-2 | 0.80 |
| 2171 | Paw-26 | 1.20 |
| 7172 | HI-3 | 1.20 |
| 6909 | Col-0 | 1.40 |
| 6944 | NFA-8 | 1.41 |
| 8310 | Hs-0 | 1.46 |
| 6973 | UII2-3 | 1.54 |
| 6976 | Uod-7 | 1.55 |
| 765 | Sus-1 | 1.58 |
| 8334 | Lu-1 | 1.70 |
| 7382 | Utrecht | 1.77 |
| 8343 | Na-1 | 1.80 |
| 7015 | Bla-1 | 1.87 |
| 7477 | WAR | 1.88 |
| 6987 | Ak-1 | 1.89 |
| 8354 | Per-1 | 1.99 |
| 6962 | Shahdar<br>a | 2.03 |
| 7309 | Po-1 | 2.04 |
| 1874 | MNF-Pot-80 | 2.30 |
| 9873 | Ndc-0 | 2.35 |
| 7268 | Np-0 | 2.37 |
| 6992 | Ang-0 | 2.43 |
| 6980 | Ws-0 | 2.61 |
| 6034 | Hov1-7 | 2.63 |
| 6979 | Wei-0 | 2.71 |
| 7004 | Bs-2 | 2.83 |
| 266 | RAN | 3.61 |
| 7418 | Zu-1 | 3.73 |
| 6284 | TV-38 | 3.99 |
| 5719 | Bur-0 | 4.34 |
| 7438 | N13 | 4.71 |
| 7404 | Wc-1 | 5.34 |
| 1552 | Sku-30 | 5.53 |
| 766 | Dja-1 | 6.37 |
| 7307 | Pn-0 | 6.40 |
| 6153 | TAA 03 | 6.75 |
| 9427 | Näs 2 | 7.50 |
| 6025 | Gro-3 | 7.59 |
| 6900 | Bil-5 | 7.68 |
| 7353 | Tha-1 | 7.74 |
| 6978 | Wa-1 | 8.26 |
| 7242 | Lo-2 | 8.40 |
| 9089 | Nar-3 | 8.89 |
| 6043 | Löv-1 | 9.22 |
| 7352 | Te-0 | 9.42 |
| 6016 | Eds-1 | 9.56 |
| 7181 | Je-0 | 9.58 |
| 9388 | Grön 14 | 9.67 |
| 6977 | Van-0 | 9.83 |
| 6235 | TOM 01 | 9.85 |
| 6188 | TDr-1 | 10.34 |
| 6177 | TÅL 03 | 10.41 |
| 9831 | Cas-0 | 10.43 |
| 6169 | TÅD 01 | 10.45 |
| 9321 | Ådal 1 | 10.48 |
| 9433 | Nyl 13 | 10.54 |
| 9363 | EdJ 2 | 10.75 |
| 9354 | Eden 15 | 10.76 |
| 8230 | Algutsru<br>m | 10.88 |
| 6917 | Fäb-2 | 12.49 |
| 8326 | Lis-1 | 12.79 |
| 6184 | TBÖ 01 | 12.85 |
| 6199 | TDr-14 | 12.94 |
| 5856 | Dör-10 | 13.12 |
| 6237 | TOM 03 | 13.47 |
| 6946 | Oy-0 | 13.78 |
| 6070 | Omn-1 | 13.99 |
| 5835 | Bil-3 | 14.05 |
| 9332 | Bar 1 | 14.66 |
| 5829 | Ale1-2 | 15.46 |
| 6214 | TFÅ 04 | 15.78 |
| 998 | Ale-Stenar-57-16 | 18.22 |
| 6009 | Eden-1 | 19.08 |
| 6231 | TNY 04 | 20.01 |

**Table S2.**
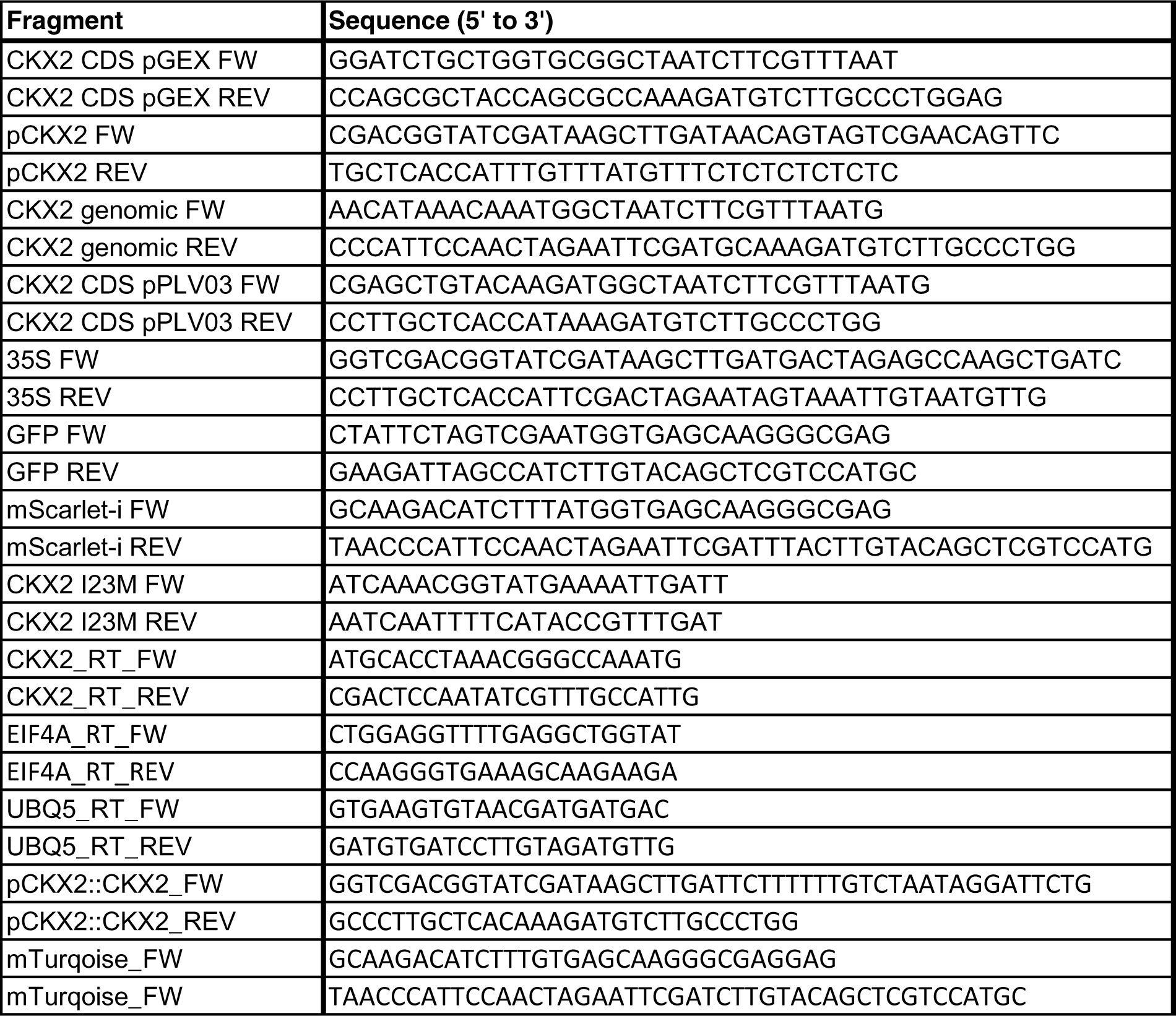
Oligonucleotides used in this study.

**Figure S1.**
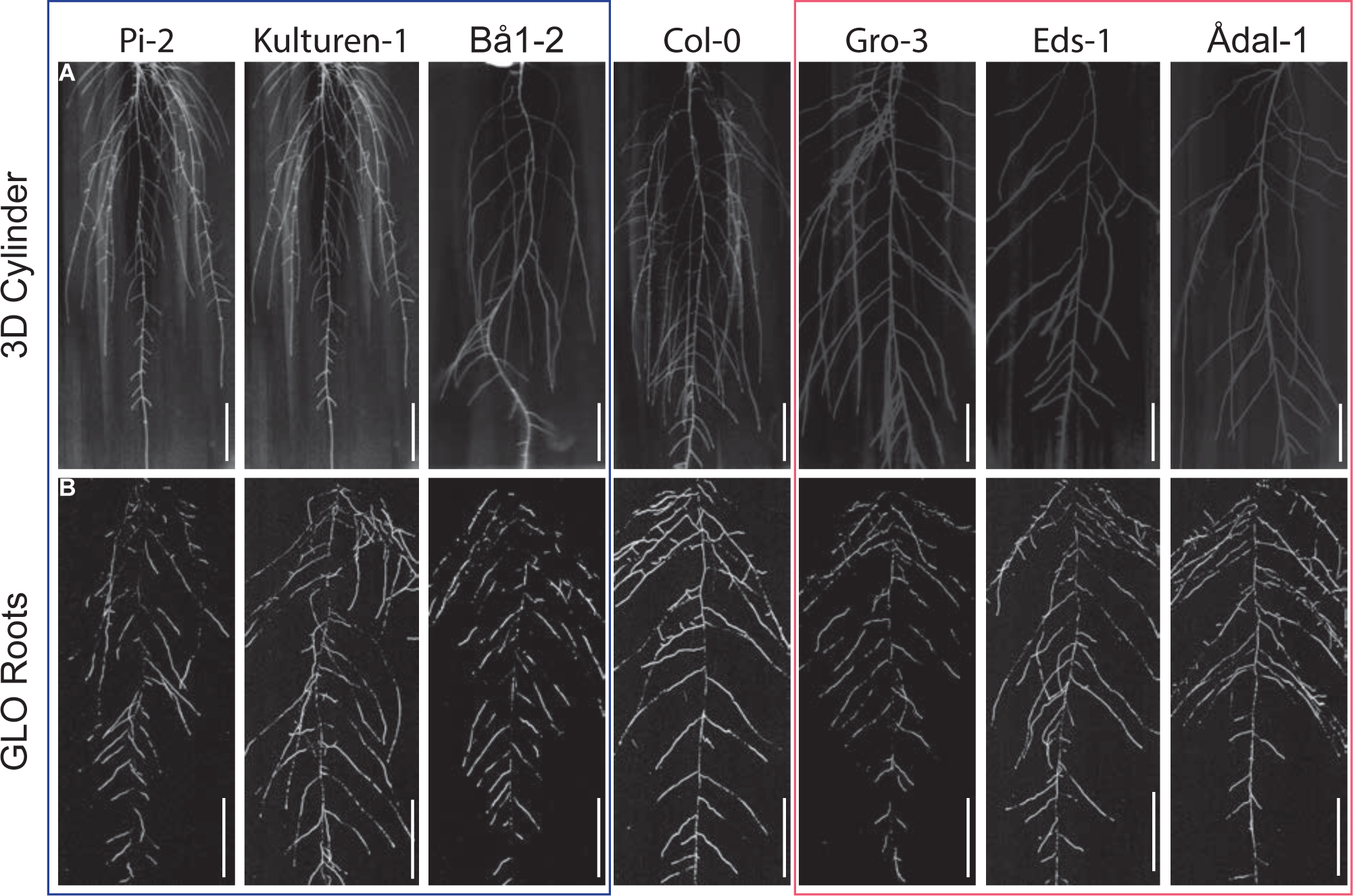
Representative root system images of selected accessions. (A) 10-day old accessions grown in 3D agar cylinders. n = 5 cylinders. Scale bars, 1 cm. (B) 20-day old accessions grown in soil. n = 5-10 plants. Scale bars, 5 cm. (A)-(B) Representative images are shown. Experiments were repeated at least three times.

**Figure S2.**
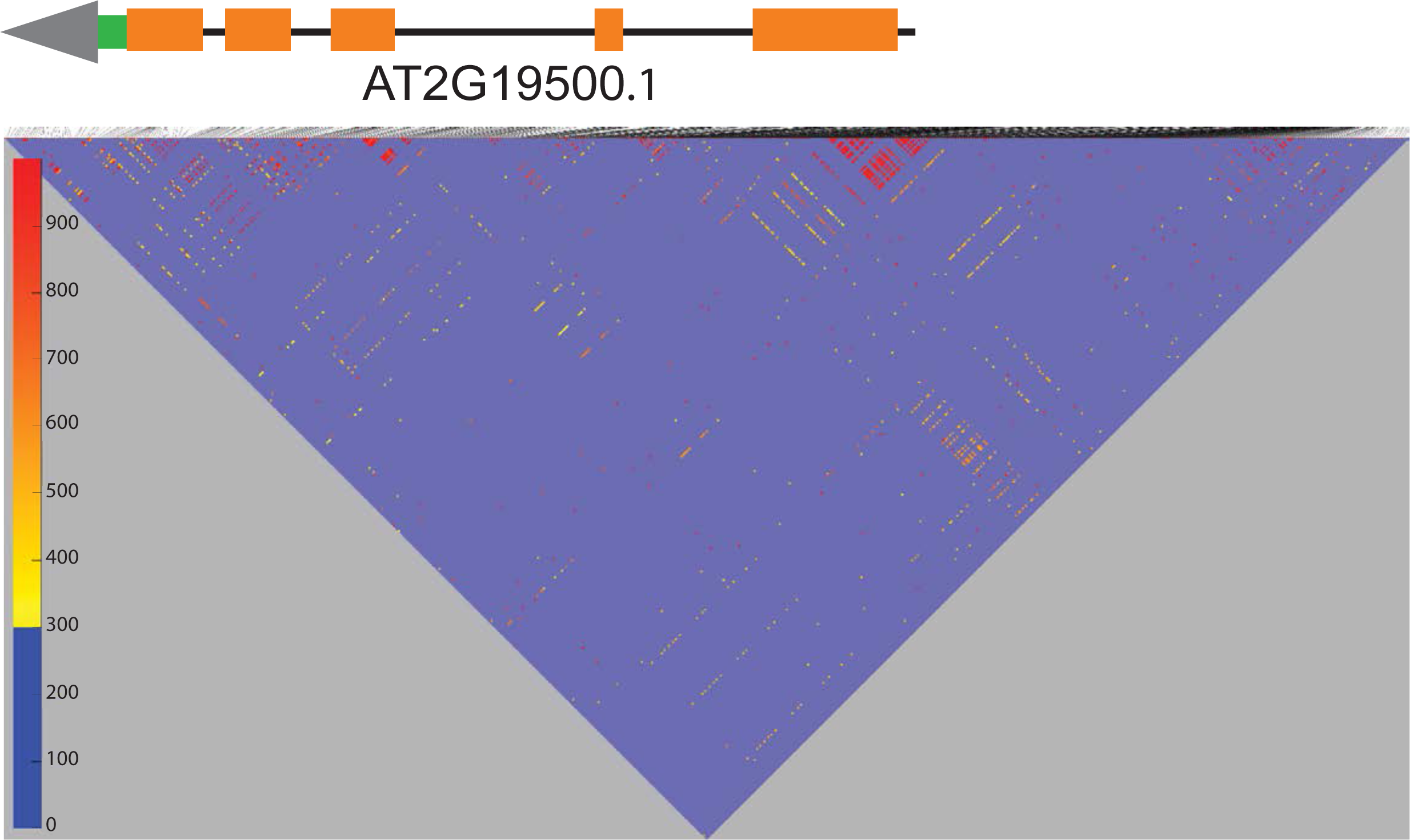
Calculation of linkage disequilibrium by pairwise comparison of 500 SNPs. r^2^ value is scaled and color-coded (blue to red) from 0 to 1 (low to high association). Underlying code can be found at github https://github.com/timeu/PyGWAS/blob/master/pygwas/core/genotype.py#L59).

**Figure S3.**
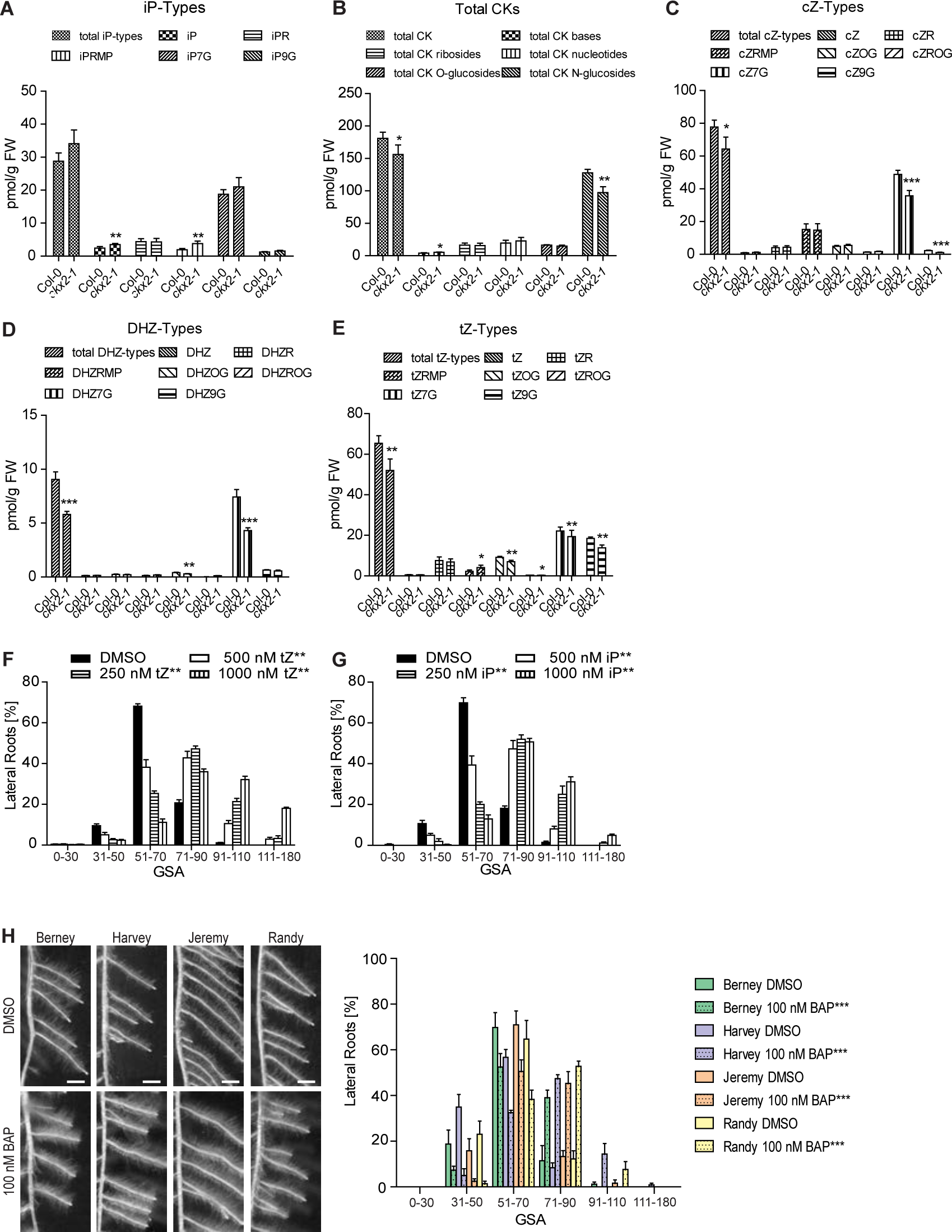
Influence of cytokinin on GSA of lateral roots. (A)-(E) Quantification of different CK forms (nucleotides (precursors), ribosides (transported forms), free bases (active forms), and O-/N-glucosides (reversible/irreversible inactivated storage forms) in Col-0 wild type and *ckx2-1* mutant roots. (A) iP-Types, (B) Total CKs, (C) cZ-Types, (D) DHZ-Types, (E) tZ-Types. (F)-(G), GSA distributions of DMSO and (F) trans-zeatin (tZ)-treated or (G) isopentenyladenine (iP)-treated Col-0 wild type seedlings. (H) Representative images and GSA distributions of four different untreated and BA-treated winter oilseed rape (Brasica napus L.) genotypes. (A)-(E) One-way ANOVA analysis P-values: * P < 0.05, ** P < 0.01, *** P < 0.001. Mean ± SD, n = 5 extractions. (F)-(H) Kolmogorov-Smirnov test P-values: * P < 0.05, ** P < 0.01, *** P > 0.001 (compared to DMSO or Col-0, respectively). Mean ± SEM, for A. thaliana: n = 5 plates (16 seedlings with 100-160 LRs per plate), for oilseed rape: n = 3 seedlings with 25-50 LRs per seedling). Experiments were repeated at least three times.

**Figure S4.**
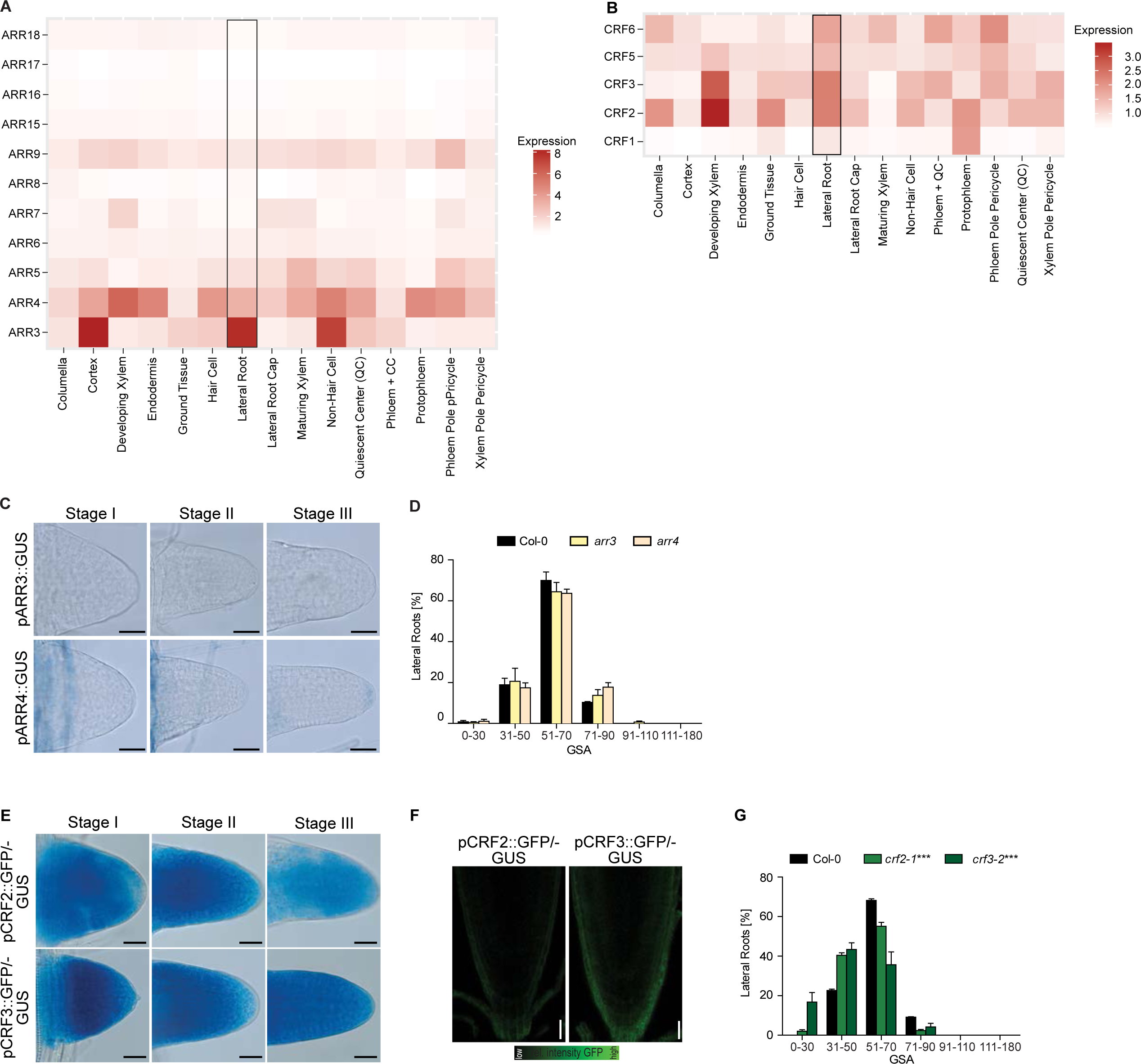
Role of type-A Arabidopsis response regulators (ARRs) and cytokinin response factors (CRFs) in the GSA establishment in LRs. (A) Expression of *ARRs* in various root tissues. Data from Brady et al., 2007. (B) Expression of *CRFs* in various root tissues. Data from Brady et al., 2007. (C) GUS staining of pARR3::GUS and pARR4::GUS. Scale bars, 25 µM. (D) GSA distribution of Col-0 wild-type, *arr3* and *arr4* mutants. Kolmogorov-Smirnov test. Mean ± SEM, n = 5 plates (16 seedlings with 100-160 LRs per plate). (E) Representative images after GUS staining of pCRF2::GFP-GUS and pCRF3::GFP-GUS in stage I-III LRs. Scale bars, 10 µm. (F) Representative images of pCRF2::GFP-GUS and pCRF3::GFP-GUS in the main root tip. Scale bars, 25 µm. (G) GSA distribution of Col-0 wild type and (A) *crf* single mutants. Kolmogorov-Smirnov test P-values: *** P < 0.001 (compared to Col-0). Mean ± SEM, n = 5 plates (16 seedlings with 100-160 LRs per plate). (C)-(G) Experiments were repeated three times.

**Figure S5.**
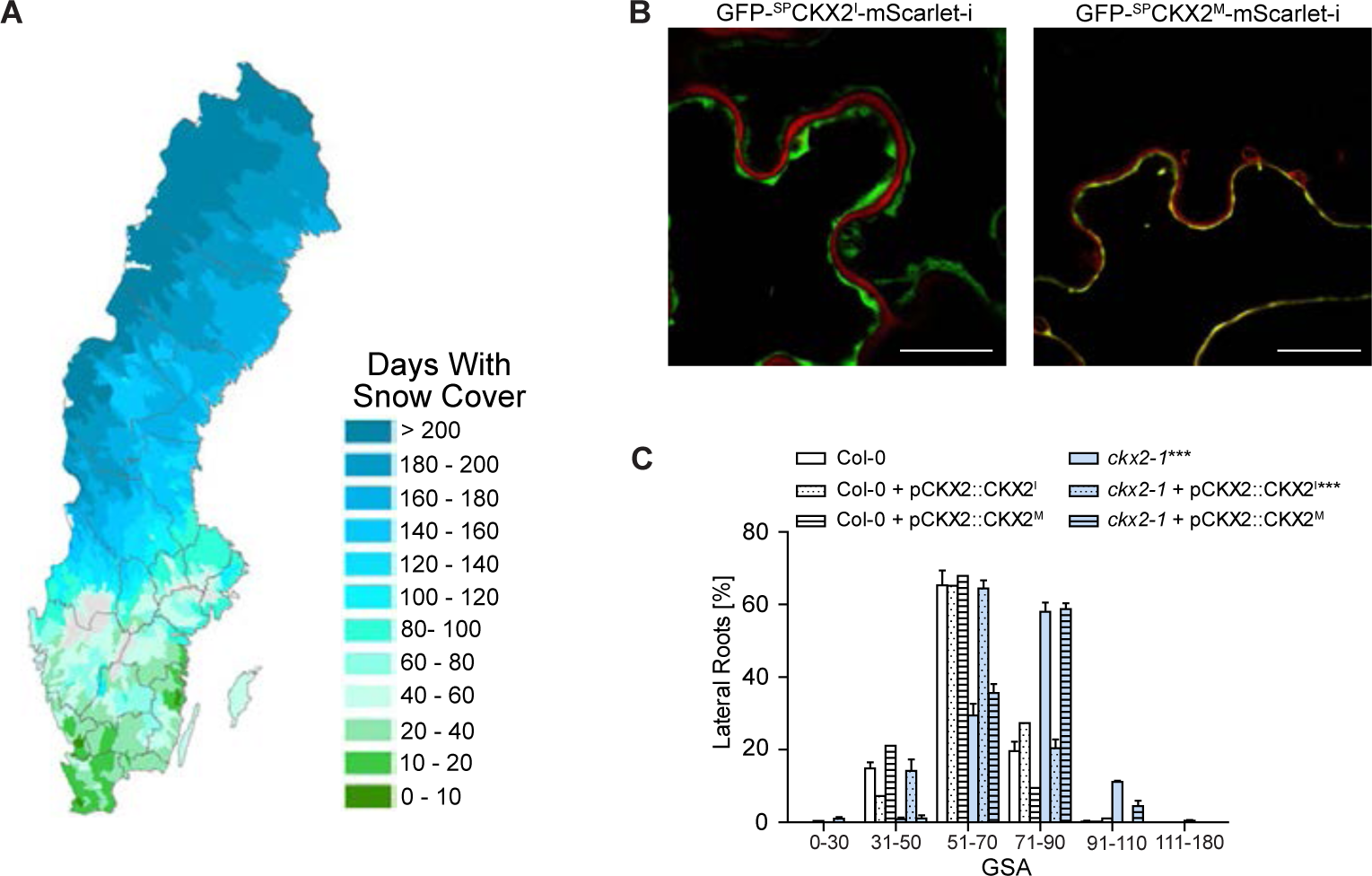
Snow cover in Sweden and characterization of CKX2^I^ and CKX2^M^. (A) Average number of days with snow cover in Sweden between 1961-1990. Source: https://bit.ly/2UmLAeT (B) Localization of GFP-^SP^CKX2^I^-mScarlet and GFP-^SP^CKX2^M^-mScarlet. Tobacco leaves were infiltrated with Agrobacterium tumefaciens containing constructs and the expression of CKX2 fluorescent protein was visualized by confocal laser scanning microscopy three days after infiltration. Scale bar, 25 µm. Representative images are shown. (C) GSA distributions of Col-0 and *ckx2-1* as well as CKX2^I^ and CKX2^M^ expressing lines in both backgrounds. Kolmogorov-Smirnov test P-value: *** P < 0.001 (compared to Col-0). Mean ± SEM, n = 5 plates (16 seedlings with 100-160 LRs per plate). (B)-(C) Experiments were repeated at least three times.

**Figure S6.**
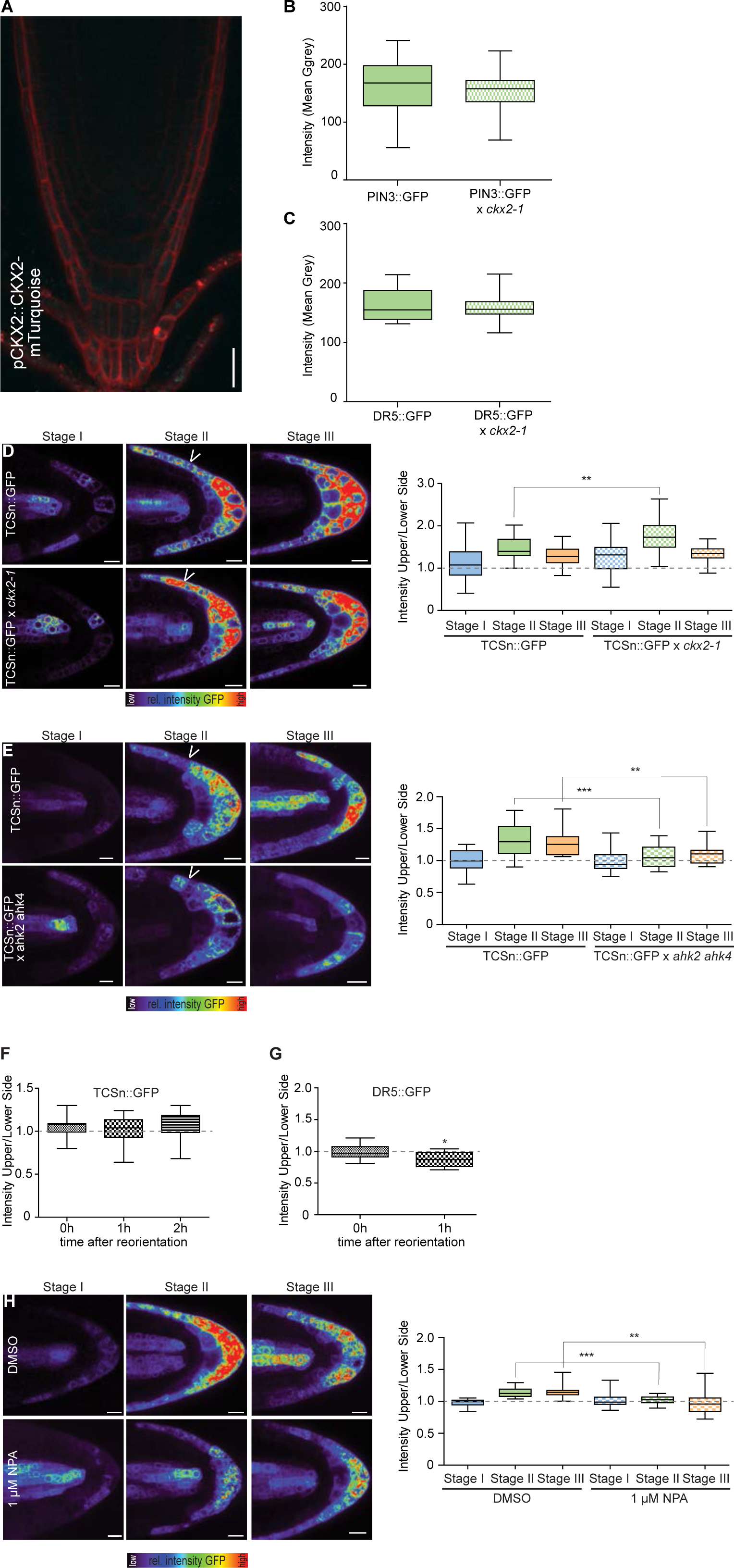
Localization of CKX2 in the main root tip and characterization of cytokinin responses in *ckx2-1* mutants. (A) Representative images of pCKX2::CKX2-mTurquoise in the main root. Propidium Iodide (PI) was used for counterstaining. Scale bar, 25 µm. (B)-(C) Signal quantification of (B) PIN3::PIN3-GFP and (C) DR5::GFP in stage II LRs. (D)-(E) Representative images and quantifications of (A) TCSn::GFP in Col-0 wild type and *ckx2-1*, (B) TCSn::GFP in Col-0 wild type and *ahk2 ahk4* in stage I – III LRs. Scale bar, 10 µm. (F) Signal quantification of TCSn::GFP in the main root after gravity stimulation. (G) Signal quantification of DR5::GFP in the main root after gravity stimulation (positive control for (F). (H) Representative images and quantifications of TCSn::GFP in Col-0 wild type after treatment with DMSO or 1 µM NPA for 24h in stage I – III LRs. Scale bar, 10 µm. (D)-(H) One-way ANOVA P-values: * P < 0.05, ** P< 0.01, *** P< 0.001. Horizontal lines show the medians; box limits indicate the 25th and 75th percentiles; whiskers extend to the min and max values. n = 10-15 individual LRs or main roots. (A)-(H)Experiments were repeated at least three times.

**Figure S7.**
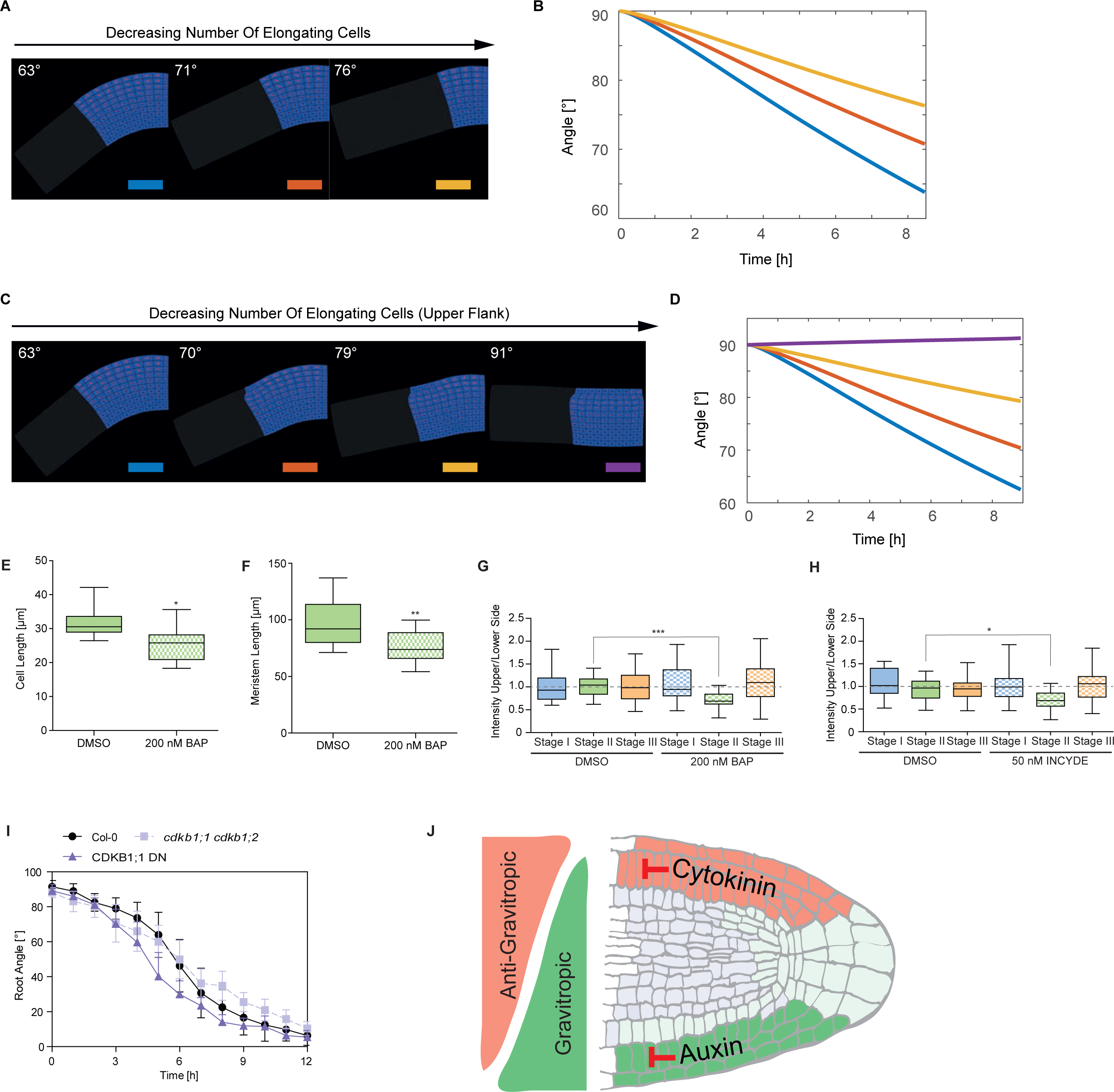
Cytokinin-dependent interference with cell division rates defines angular growth of lateral roots. (A) Computer simulations displaying that the number of elongating cells impacts on GSA angle in the proportional manner. (B) Time evolution of set-point angle corresponding to (A). (C) Decreasing number of elongating cells on the upper half part of the LR elongation zone. Note the gradual increase of set-point angle with decreasing number of elongating cells. (D) Time evolution of set-point angle corresponding to (C). Colour of curves matches simulation with the colour bar shown in (A) and (C). Simulations represent LR status after 8h of dynamic elongation. (E)-(F) Quantification of (E) first two elongated cells of stage II lateral roots and (F) lateral root meristem after treatment with DMSO or 200 nM BAP for 24h. (G)-(H) Quantification of CycB1;1::GUS after treatment with (G) DMSO or 200 nM BAP or (H) DMSO or 50 nM INCYDE for 24h. (E)-(H) One-way ANOVA P-values: * P < 0.05, *** P < 0.001. Horizontal lines show the medians; box limits indicate the 25th and 75th percentiles; whiskers extend to the min and max values. n = 10-15 individual LRs. (I) Kinetics of gravitropic response of dark grown Col-0, *cdkb1;1 ckdb1;2* and CDKB1;1 DN. Mean ± SD, n = 10-15 individual roots. (J) Schematic model depicts spatially defined gravitropic and anti-gravitropic hormonal cues at opposing organ flanks. Cytokinin signalling functions as an anti-gravitropic growth regulator at the upper side and thereby counterbalances auxin-dependent gravitropic growth of lateral roots. (A)-(I) Experiments were repeated at least three times.

